# Socio-emotional difficulties observed in alexithymia reflect altered interactions of the semantic and monoaminergic neuromodulatory brain networks

**DOI:** 10.1101/2025.05.23.655721

**Authors:** Marcos Ibáñez Montolio, Maya Visser, Michal Rafal Zareba

## Abstract

Alexithymia is a multidimensional construct characterized by difficulties in identifying and describing feelings and reduced ability to engage in abstract thinking. Although often co-occurring with other psychological and neurodevelopmental conditions such as anxiety, depression and autism spectrum disorders, alexithymia is believed to be associated with unique alterations within the socio-emotional brain networks. With the semantic and neuromodulatory brainstem systems playing a key role in social and affective cognition, the current work aimed to study their contributions to alexithymia in unprecedented detail. First, we attempted to identify resting-state functional connectivity patterns of the social semantic hubs (superior anterior temporal lobe) and monoamine-producing regions (dorsal raphe, ventral tegmental area and locus coeruleus) linked to each alexithymia domain. Secondly, by deploying tractography and graph analysis of the associated structural network, we intended to identify their potential anatomical correlates. Alexithymia was strongly associated with dysconnectivity within the semantic network, and altered functional connectivity between the neuromodulatory brainstem regions and cortical areas crucial for social cognition and emotion regulation, including medial prefrontal cortex and inferior parietal lobule. On the anatomical level, these findings were paralleled by negative links with network modularity, suggestive of less specialised neural processing, and decreased clustering coefficient of the semantic node in the left posterior middle temporal gyrus. Despite observing associations with trait-anxiety and emotion suppression for some of the highlighted findings, these phenomena did not mediate the effects of alexithymia. Therefore, the current work highlights the existence of functional and structural alterations within socio-emotional networks as neural markers of alexithymia.

## 1. Introduction

Being able to understand and describe one’s feelings, as well as infer the emotional state of others, are essential components of our everyday functioning. These cognitive processes not only form the building blocks of all social interactions but are also key predeterminants for successful emotion regulation. As such, unsurprisingly, deficits in these capabilities have adverse consequences, affecting one’s emotional well-being as well as their social life. The aforementioned “lack of words for emotions” is what Sifneos (1973) coined as “alexithymia”, a multifaceted and multidimensional construct that has been expanded over the years to include, between others, the difficulty to identify and describe one’s, or other people’s, feelings, the difficulty in distinguishing feelings from bodily sensations, a diminution of fantasy, and a decreased ability to engage in abstract thinking (De Berardis et al., 2020). Regardless of the ongoing debate on its factor structure (Goerlich, 2018), the most extensively used measure for alexithymia is the TAS-20 (20 item Toronto Alexithymia Scale; Bagby et al., 1994), a questionnaire with a three-dimensional structure assessing: difficulty identifying feelings (DIF), difficulty describing feelings (DDF) and externally oriented thinking (EOT), that is, a strong preference to attend to external objects, people, and environmental events rather than examining own feelings (Jakobson et al., 2024).

The co-occurrence of alexithymia-related deficits with a multitude of psychological conditions such as anxiety, depression and autism spectrum disorders (ASD), have led many to regard alexithymia and its effects as merely derived symptoms (Gaigg, 2012; Haviland et al., 1988; Quattrocki & Friston, 2014; Zeitlin & McNally, 1993). Such a simplification may stem from the strong link between emotional regulation and those psychological disorders (Clark & Watson, 1991; Cuve et al., 2022; Hofmann et al., 2012). For some individuals, the lack of awareness of emotions present in alexithymia may lead to affective dysregulation (Taylor et al., 1989) due to the regulation process being blocked as a direct consequence of not being able to access their adaptive informational meaning (Taylor et al., 2016). Therefore, contrary to the classic literature, alexithymia may facilitate the development of mood and anxiety symptomatology. This theory is supported by the greater prevalence of maladaptive emotion regulation strategies (i.e. expressive suppression, cognitive avoidance, catastrophizing, or blaming others) in individuals with high alexithymia scores (Abdi et al., 2023; Luo et al., 2022; Walker et al., 2011).

On the same note, the current body of research also points towards symptomatological and neuroanatomical independence of alexithymia (Bernhardt et al., 2014; Kinnaird et al., 2019; Marchesi et al., 2000). The described facets of alexithymia rely on the neural network involved in socio-emotional cognition. However, the patterns of involvement within this network will most likely differ depending on the exact construct (e.g., identifying vs describing feelings) and will be different from those related to often co-occurring diagnostics (e.g., anxiety or ASD). Nevertheless, individual symptomatology is frequently shared between alexithymia and said conditions. For example, a lack of emotional awareness in people with high alexithymia scores has been linked to difficulties in interpreting social situations (Di Tella et al., 2020), a symptom commonly associated with ASD (Gaigg, 2012). Similarly, alexithymia has been associated with reduced abilities in perspective-taking and empathic concern (Alkan Härtwig et al., 2020; Moriguchi et al., 2007), which are key components of guilt feelings (Martinez et al., 2014). As patients with anxiety, depression and ASD cope with decreased empathic feelings towards one-self (Cai et al., 2023; Garnefski et al., 2002), alexithymia could potentially be an explanatory factor behind these shared characteristics.

A key region for processing such social information is the superior anterior temporal lobe (ATL), believed to be the locus for social semantics in the brain (Binney & Ramsey, 2020; Olson et al., 2013). Aberrant functional connectivity of this area with fronto-limbic regions has been associated with guilt processing in depression (Zahn et al., 2009; Lythe et al., 2015) and subclinical anxiety (Zareba et al., 2024b), underlying the important role of this region in emotional processing and regulation across mental health conditions. It has also been shown that the activity in the superior ATL is linked to interpreting social interactions (Olson et al., 2013) and understanding moral concepts (Zahn et al., 2007). However, little is known about the influence that alexithymia’s dimensions have upon the social conceptual (i.e., ATL) and emotional (fronto-limbic) networks. Furthermore, most established theories posit the monoaminergic serotonin, dopamine and noradrenaline systems as the key elements enabling socially-driven emotions and cognitions (Avery & Krichmar, 2017; Eslinger et al., 2021). Therefore, investigating how alexithymia influences the functioning of the ATL and especially its interactions with the aforementioned fronto-limbic and neuromodulatory circuitry could greatly enhance our understanding of social emotions, possibly opening new therapeutic avenues.

In order to fill this crucial gap of knowledge, as the first goal, the current study assessed how resting-state activity and functional connectivity within these brain networks relate to the distinct facets of alexithymia. In specific, we focused on the bilateral ATL, involved in social cognition, and on the neuromodulatory circuitry, i.e. the dorsal raphe (DR), ventral tegmental area (VTA) and bilateral locus coeruleus (LC), representing the monoaminergic serotonin, dopamine, and noradrenaline systems, respectively.

Prior literature regarding the involvement of brainstem modulatory systems in alexithymia has been dominated by genetic studies (Li et al., 2020; Terock et al., 2021; Yang et al., 2019), while investigations on the neural level remain scarce. Despite some studies reporting associations between alexithymia and the genotypes of COMT and 5-HTTLPR, a recent meta-analysis by Yang et al. (2019) failed to find consistent effects, although this lack of consistency related to the monoaminergic system can also be observed in other conditions where emotional regulation plays an important role, such as high-trait anxiety (Berry et al., 2019; Zalachoras et al., 2022). Similarly, prior research has not focused on the functional characteristics of the ATL. Although labelling of others’ emotions is paralleled in highly alexithymic individuals by increased activity in the social cognition network, including left inferior frontal gyrus, precuneus and the right temporal lobe (Alkan Härtwig et al., 2020), this finding is not easily translatable to the resting-state paradigm, as passive cognitive processes taking place during rest (such as “mind wandering”) may differ in their demands from more complex social cognitions. Therefore, our investigation is meant to weigh additional arguments into the ongoing debate on how the activity of neuromodulatory areas and the ATL is associated with alexithymia.

Additionally, recent evidence extracted from animal studies on rodents (Ren et al., 2018) has shown that the manipulation of distinct connections on the same neuromodulatory region can exert opposite effects on behaviour. As such, we expected to observe both negative and positive correlations of alexithymia with the resting-state connectivity of the aforementioned monoaminergic areas. On the other hand, a pilot study deploying neurofeedback (Dobrushina et al., 2022) reported that decreases in alexithymia scores correlated with increased connectivity between the right temporal pole and the visual cortices involved in semantic processing, as well as with higher connectivity within a network involving socio-emotional processing areas such as the anterior insula and the temporoparietal junction. These findings suggest the existence of deficits in automatic socio-emotional processing that disappear when a more prolonged conscious effort to engage is made (Ihme et al., 2014). With resting-state coupling reflecting to a greater extent the habitual co-activation patterns between the brain areas (Guerra-Carrillo et al., 2014), we expected alexithymia to be negatively associated with the functional connectivity between the superior ATL (sATL) and the previously discussed circuitry.

The second goal of the study was to examine whether the neural architecture underlying the functional connections highlighted in the aforementioned analysis was related to both alexithymia and the functional findings. On one hand, taking a straightforward approach, this objective was achieved through performing tractography between the pairs of brain regions from the identified functional connectivity patterns. On the other hand, understanding that not all functionally connected areas will be directly linked by white matter (WM) tracts, we constructed in each subject a structural network encompassing the anatomical connections between all the regions from the functional analysis, and used graph theory to derive local and global network parameters. As demonstrated in a recent study by Sarwar et al. (2021), there exists a close relationship between the characteristics of anatomical and functional connections, as well as the associated cognitive functions. As such, we hypothesised that for the functional connectivity patterns with an existing direct anatomical link, the structural properties of the associated WM tracts will be related to both alexithymia and the strength of the functional coupling between the brain areas. Furthermore, as previous studies using graph theory have found correlations for the nodal and global network parameters with emotional processing (Zareba et al., 2024b) and awareness (Smith et al., 2018), we expected to observe similar links with alexithymia in our study.

Lastly, we investigated whether the alexithymia-related effects in the highlighted circuitry were mediated by the external factors previously reported to be closely associated with this condition, i.e. trait-anxiety and suppression/reappraisal emotional regulation strategies (Berthoz et al., 1999; Luo et al., 2022; Walker et al., 2011). We hypothesised that neither will show a mediating effect, adding to the current body of knowledge pointing towards the delineation of alexithymia as an independent construct (Bernhardt et al., 2014; Kinnaird et al., 2019; Marchesi et al., 2000).

## 2. Materials and methods

### 2.1. Participants

The sample employed in this study was assembled from the anonymised magnetic resonance imaging (MRI) and behavioural data provided by the *Max Planck Institute Mind-Brain-Body Dataset* (Babayan et al., 2019), an open access dataset compliant with the Declaration of Helsinki’s ethical standards. The inclusion criteria to be met by the participants were: being 20 to 35 years old, right handedness, having at least achieved secondary-level education, and reporting no concurrent drug consumption nor past neurological, psychiatric or substance use disorders. The sample for the functional analysis consisted of 60 participants (20 females). One of the subjects (female, 20-25 age range) was excluded from the diffusion MRI (dMRI) analysis due to incomplete data (lack of diffusion-weighted fieldmaps).

### 2.2. Behavioural data

Alexithymia and its various facets were assessed through the use of the Toronto Alexithymia Scale (TAS-26; Bagby et al., 1994; Kupfer et al., 2001) due to the fact that the revised version of this questionnaire (TAS-20) displays low reliability concerning the EOT scale in the German version (Kupfer et al., 2001). The tool consists of 26 items scored on a 1 to 5 Likert scale (strongly disagree - strongly agree) divided into DIF, DDF and EOT subscales. An overall score can be obtained by adding the scores from these subscales. Additionally, the Emotion Regulation Questionnaire (ERQ; Abler & Kessler, 2009; Gross & John, 2003), consisting of 10 items scored on a 1 to 7 Likert scale (strongly disagree - strongly agree), was employed to assess the habitual use of cognitive reappraisal and expressive suppression strategies for emotion regulation. Trait-anxiety was assessed using the State-Trait-Anxiety Inventory (STAI; Laux et al., 1981; Spielberger, 1989; Spielberger et al., 1970), a 20-item questionnaire rated on a Likert scale from 1 to 4 (almost never - almost always). Lastly, semantic cognition-related scores were extracted from the Regensburger Wortflüssigkeits-Test (RWT; Aschenbrenner et al., 2000) and the Wortschatztest (WST; Schmidt & Metzler, 1992). In specific, RWT-1 measures the total number of words starting with “s” said by the participant in 1 minute, while RWT-13 measures the total number of animals said in the same timescale. Lastly, WST-1 collects the total number of real words recognized from 42 word sets that also include 5 pseudo-words. A demographic and behavioural summary of the sample is provided in ***Table 1***.

**Table 1.** Demographic and behavioural data.

|  |  |  |
| --- | --- | --- |
| Gender | Male | 40 (66.67%) |
|  | Female | 20 (33.33%) |
| Age range | 20-25 | 30 (50%) |
|  | 25-30 | 25 (41.67%) |
|  | 30-35 | 5 (8.33%) |
| TAS (Mean ± SD) | Difficulty Identifying Feelings | 12.9 ± 4.29 |
| | Difficulty Describing Feelings | 14.35 ± 4.21 |
| | Externally Oriented Thinking | 14.08 ± 3.58 |
| | Overall Score | 41.33 ± 8.43 |
| ERQ (Mean ± SD) | Suppression | 3.54 ± 1.08 |
| | Reappraisal | 4.58 ± 0.84 |
| STAI (Mean ± SD) | Trait-Anxiety | 36.95 ± 8.04 |
| RWT (Mean ± SD) | Phonetic Fluency (RWT-1) | 15.1 ± 3.45 |
| | Semantic Fluency (RWT-13) | 25.28 ± 5.60 |
| WST (Mean ± SD) | Real Word Recognition (WST-1) | 33.32 ± 2.72 |

**TAS:** Toronto Alexithymia Scale; **ERQ:** Emotion Regulation Questionnaire; **STAI:** State-Trait-Anxiety Inventory; **RWT:** Regensburger Wortflüssigkeits-Test; **WST:** Wortschatztest; **SD:** Standard Deviation

### 2.3. MRI data acquisition

Acquisitions were performed on a 3 Tesla scanner (MAGNETOM Verio, Siemens Healthcare GmbH, Erlangen, Germany) equipped with a 32-channel head coil. Structural images were obtained with an MPRAGE sequence (176 sagittal slices; 1 mm isotropic voxel size; TR = 5000 ms; TE = 2.92 ms; flip angle 1/flip angle 2 = 4°/5°; GRAPPA acceleration factor 3). For the T2*-weighted resting-state functional sequence, participants were required to focus their eyes on a low-contrast fixation cross. A gradient echo planar multiband imaging sequence was employed to attain the data (64 axial slices acquired in interleaved order; 2.3 mm isotropic voxel size; TR = 1400 ms; TE = 30 ms; flip angle = 69°; multiband acceleration factor = 4). 657 volumes were obtained, making the total scan duration 15 min 30 s. In addition, the whole-brain high angular resolution diffusion-weighted images were collected axially (88 slices in the interleaved order) with a 1.7-mm isotropic resolution (60 diffusion-encoding gradient directions, *b*-value of 1000s/mm^2^, 7 *b*_0_ images, TR = 7000 ms, TE = 80 ms, FA = 90°, GRAPPA acceleration factor = 2, 32 reference lines, multiband acceleration factor 2).

### 2.4. Functional analysis

#### 2.4.1. Data preprocessing and (f)ALFF computation

As per the methodology followed in previous studies conducted by our research group (Zareba et al., 2024a), the first 5 volumes for the functional MRI (fMRI) data were discarded in order to guarantee signal stabilisation. This functional and anatomical data was then fed into the fMRIPrep 23.0.0 (Esteban et al., 2019), a data preprocessing pipeline based on DRNipype 1.8.5 (Gorgolewski et al., 2011). As a summary of the undergone steps, the anatomical images were skull-stripped, segmented and normalized to the MNI (Montreal Neurological Institute) space. Similarly, functional data underwent motion, susceptibility distortion and slice-timing correction, was coregistered to the anatomical image and normalized.

Subsequent preprocessing steps and computation of Amplitude of Low-Frequency Fluctuations (ALFF) and fractional ALFF (fALFF) were conducted in AFNI (Cox, 1996). The fMRI volumes were smoothed with a 4 mm Gaussian filter, detrended, denoised and band-pass filtered in the 0.01-0.1 frequency range. The denoising regressors, which included 6 motion parameters, their derivatives, and component-based physiological regressors (aCompCor; Behzadi et al., 2007) for cerebrospinal fluid (CSF), were extracted from the fMRIPrep output.

As recommended by Woletz et al. (2019), the fMRI data was not subjected to polynomial detrending for fALFF calculations. ALFF was computed as the total signal power within the 0.01-0.1 Hz frequency range, and fALFF as a ratio of the signal power within this range divided by the signal power of the entire frequency spectrum. Both metrics were mean-normalized within each subject preceding the group analysis.

Due to the inherent differences between ALFF and fALFF, with ALFF being more sensitive to physiological noise (Zou et al., 2008), but showing greater test-retest reliability in grey matter regions than fALFF (Zuo et al., 2010), it was determined that this investigation would benefit from including both metrics.

#### 2.4.2. Regions of interest

The subcortical regions of interest (ROIs), i.e. DR, LC and VTA, were delineated in MNI space using publicly available resources. Specifically, in their original articles (Believau et al., 2015; Betts et al., 2017), DR and bilateral LC seeds were defined in a data-driven manner using 5-HTT transporter binding maps and T1-weighted FLASH sequence, respectively. In turn, VTA seed was originally hand-drawn by experts using the established anatomical landmarks (Murty et al., 2014). Additionally, we defined two ATL ROIs for each hemisphere using the rostral superior temporal gyrus subdivisions (i.e. STG0 and STG2) from Hoffman & Lambon Ralph’s work (2018). See ***Supplementary Figure 1*** for visual representation.

#### 2.4.3. Group-level resting-state activity analysis

The ALFF and fALFF values were obtained by extracting the mean for every ROI in each of the subjects. Linear regression models were employed to calculate the associations between the amplitude of resting-state neural activity and alexithymia scores using R (version 4.3.0; R Core Team, 2021). For each ROI, the ALFF and fALFF values were designated as the dependent variable. Gender, age and each of the TAS-26 scales were used as the regressors. Multiple comparisons correction was established accounting for false-discovery rate (FDR < 0.05).

#### 2.4.4. Group-level whole-brain connectivity analysis

The seed time courses of the blood-oxygen-level-dependent (BOLD) signal were calculated by averaging the time-series from all voxels in each ROI, and were subsequently correlated through Pearson’s method with the time courses of all the other voxels in the brain to create individual whole-brain functional connectivity maps. The distribution of the correlation coefficients was normalized using Fisher’s z transform.

AFNI’s 3dMVM program (Chen et al., 2014) was used to conduct analysis of covariance (ANCOVA) to assess the associations of each alexithymia facet with each ROI’s whole-brain functional connectivity pattern. In this regard, gender and age were designated as covariates. Voxel-level thresholding was set at a p < 0.001 level, while cluster-level family-wise error rate correction (FWE < 0.05) was conducted to account for multiple comparisons. Mean functional connectivity values between surviving clusters and their pertaining ROIs were extracted for each subject and subsequently correlated with RWT and WST semantic scores to test whether the highlighted neural coupling patterns were specific to alexithymia, or reflected interindividual variability in more generalised semantic processing. FDR (< 0.05) correction method for multiple comparisons was applied.

### 2.5. Diffusion analysis

#### 2.5.1. Data preprocessing

Following the methodology employed in prior studies published by our research group (Zareba et al., 2024b), preprocessing of T1-weighted data was carried out using the Computational Anatomy Toolbox (CAT12; Gaser et al., 2024). The individual anatomical images were skull-stripped and segmented into grey matter (GM), WM and CSF masks, and were subsequently used for delineation and coregistration of ROIs to other MRI modalities. AFNI (Cox, 1996) was used to create the skull-stripped anatomical image based on the whole-brain mask obtained using CAT12, as this tool does not automatically output this type of images.

Preprocessing of the dMRI data was performed using the MRtrix3 software (Tournier et al., 2019). Denoising and unringing of the images was followed by distortion, eddy currents and motion correction using the dwifslprepoc wrap-up rooted in the functionalities offered by the FMRIB Software Library (FSL; Jenkinson et al., 2012). The diffusion images derived from this procedure were then bias-field corrected. The dhollander algorithm (Dhollander et al., 2019) was applied to estimate the response functions for WM, GM and CSF, which were employed as input for the single shell 3 tissue-constrained spherical deconvolution carried out using MRtrix3Tissue (https://3Tissue.github.io), a fork of MRtrix3 (Tournier et al., 2019). Intensity normalization was subsequently accompanied by anatomically constrained tractography (ACT; Smith et al., 2012) based on the 5-tissue image outputted by the *5ttgen* function of the fsl algorithm. Compared to previous tractography algorithms, ACT improves the biological accuracy of the reconstructions by seeding from the GM-WM interface and terminating the connections upon entering cortical or subcortical GM, therefore reducing the number of known false positives. 10 million streamlines were generated for each subject using the improved second-order integration over fiber orientation distributions algorithm (iFOD2; Tournier et al., 2010). Default settings were applied except for the maximal track length, set to 100mm, and the cut-off value, set to 0.06. Lastly, the SIFT2 algorithm (Smith et al., 2015a) was employed to refine the generated streamlines in order to enable the use of the number of streamlines as a valid index of structural connection density.

#### 2.5.2. Region of interests

ATL, DR, bilateral LC, VTA and the surviving clusters from the group-level functional analysis were selected for the diffusion analysis for a total of 12 ROIs (8 cortical and 4 subcortical). All the following steps were conducted in AFNI (Cox, 1996).

Cortical ROIs were delineated in the individual brains by warping them from the MNI space to native diffusion space. These ROIs were then manually edited in the anatomical resolution, ensuring that no voxels were located outside the GM, and were subsequently resampled to the diffusion-weighted imaging (DWI) resolution. Due to the distortion that the warping process would create for the subcortical ROIs, these were instead drawn manually in the anatomical resolution as per Levinson et al. (2023), and resampled to the DWI resolution. Visual inspection and further correction were implemented to ensure proper mapping of the ROIs once in the DWI resolution prior to combining them into individual atlases for each subject. These atlases were then used to create structural connectivity matrices and extract the mean fractional anisotropy (FA) and apparent diffusion coefficients (ADC) values for all the connections in each participant using the tck2connectome function (Smith et al., 2015b) in MRtrix3 (Tournier et al., 2019).

#### 2.5.3. Group-level analysis

Structural connectivity matrices were binarized and combined to calculate the number of participants in which each specific connection was present (see ***Figure 1***). Mean number of streamlines, FA and ADC values were extracted for the structural connections between pairs of regions stemming from the functional connectivity analysis, on condition that a direct anatomical link was found in more than one third of the sample. Linear regression models were then employed to calculate the associations between these connection-specific values, designated as the dependent variables, and alexithymia scores using R (version 4.3.0; R Core Team, 2021). Gender, age and TAS-26 subscales values were employed as regressors. False-discovery rate (FDR < 0.05) was used to correct for multiple comparisons. Linear regression models were then employed to determine the association between functional and structural connectivity values for each of these specific connections in R (version 4.3.0; R Core Team, 2021). In this regard, functional connectivity was set as the dependent variable while the number of streamlines, FA and ADC values were used as regressors.

**Figure 1.**
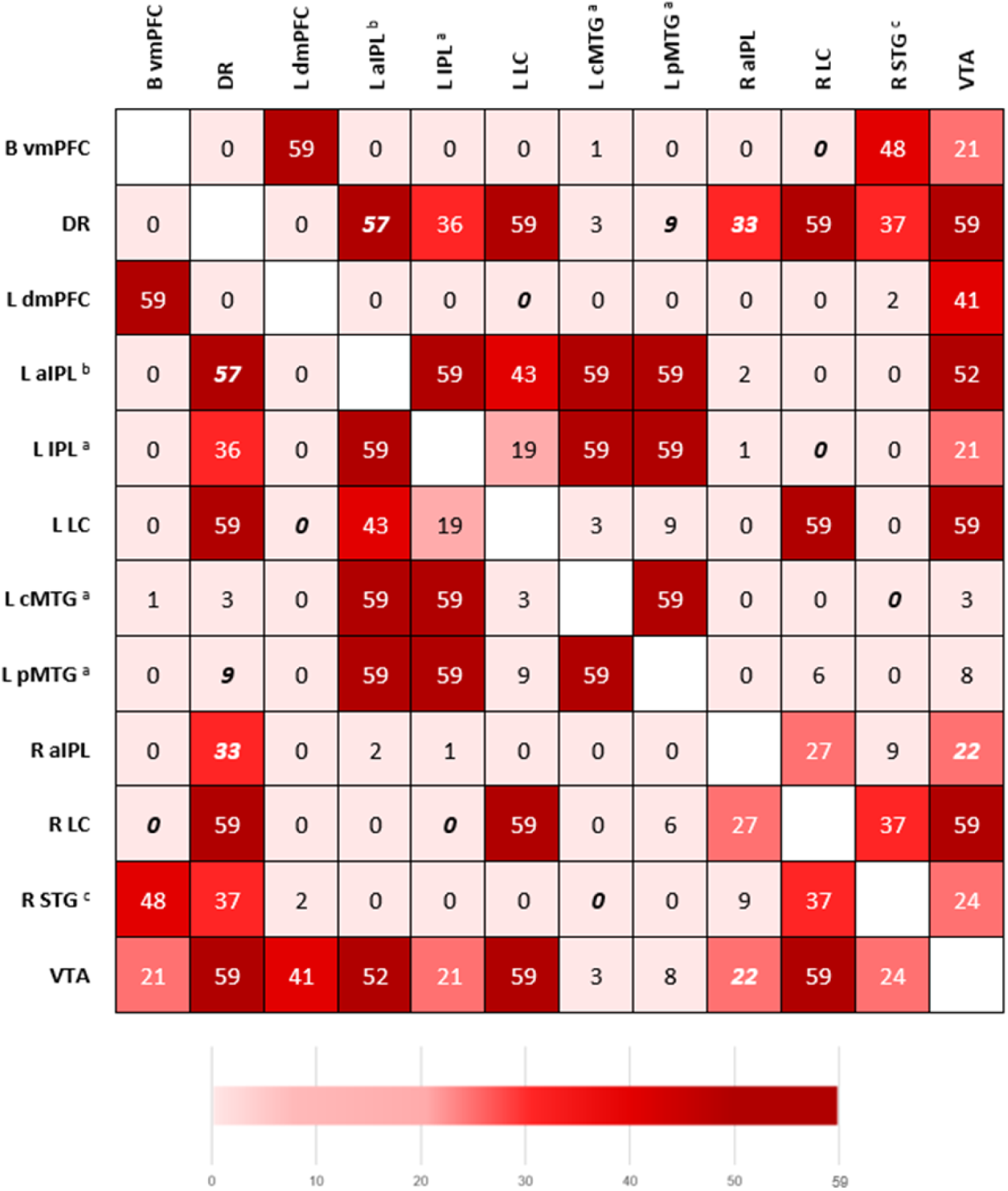
Connectivity matrix with the number of participants in which each structural connection was present. Structural connections appearing in the functional connectivity analysis are shown in italics. **L**: Left; **R**: Right; **B**: Bilateral; **vmPFC**: ventromedial Prefrontal Cortex; **DR**: Dorsal Raphe; **dmPFC**: dorsomedial Prefrontal Cortex; **IPL**: Inferior Parietal Lobule; **aIPL**: anterior IPL; **LC**: Locus Coeruleus; **cMTG**: central Middle Temporal Gyrus; **pMTG**: posterior MTG; **STG**: Superior Temporal Gyrus; **VTA**: Ventral Tegmental Area. ^a^ Peak activity coordinates, MNI: **L IPL** (−46, −66, 46); L cMTG (−60, −61, 0); **L pMTG** (−64, −25, −5). ^b^ Center of mass coordinates, MNI: **L aIPL** (−56, −38, 51). **L aIPL** includes two overlapping areas from the functional analysis: **L IPL / PCG** (Postcentral Gyrus) (peak activity coordinates: −55, −32, 57) and **L IPL** (peak activity coordinates: −57, −41, 50). ^c^ This ROI pertains to the **R STG0** subdivision as described in the work by Hoffman and Lambon-Ralph (2018).

#### 2.5.4. Graph theory analysis

Using the igraph package (Csardi & Nepusz, 2006) in R, weighted undirected graphs were created employing the streamline matrices obtained during the previous analysis. Given the small size of the network, no density thresholding was applied, i.e. all the connections were included in the calculations, regardless of their strength. Modularity (Leiden algorithm; Traag et al., 2019), global efficiency and mean clustering coefficient were calculated as the global network parameters. Weighted clustering coefficient and betweenness centrality were extracted on the nodal level. Definitions for these measures are provided in the supplementary material (***Supplementary Table 1***) (Barrat et al., 2004; Freeman, 1979; Latora & Marchiori, 2001; Traag et al., 2019).

Given the non-Gaussian distribution of multiple graph metrics, permutation-based linear regression models (10000 permutations) were used to test the associations between alexithymia scores and the graph indices using the permuco package (Frossard & Renaud, 2021). The graph measures were set as dependent variables, while gender, age and TAS-26 subscales scores were designated as regressors. Results were then corrected for multiple comparisons using false-discovery rate method (FDR < 0.05), and correlated with the motion parameters and indices of general semantic cognition (i.e. RWT and WST) to assess their possible confounding effects.

### 2.6. Mediation analysis

In order to examine if trait-anxiety and emotion regulation strategies (reappraisal and suppression) mediate the relationship between alexithymia and the neuroimaging findings, firstly we used linear regression models to calculate the associations of these variables with the TAS-26 scores and the neuroimaging findings. Gender and age were added as regressors in all models. As we tested associations with multiple neuroimaging patterns, FDR < 0.05 was used to control for multiple comparisons. Following the suggestion of Baron & Kenny (1986), only the resulting significant findings were employed in the elaboration of the mediation models, as no mediation effect would be found between unrelated variables.

Mediation models were subsequently elaborated using R’s mediation package (Tingley et al., 2014). TAS-26 subscales were set as the independent variables, trait-anxiety, suppression and reappraisal were individually selected as mediators where appropriate, while the neuroimaging findings were included as the dependent variables.

## 3. Results

### 3.1. Resting-state amplitude and functional connectivity of the socio-emotional brain regions

We observed a nominally significant negative association between DIF and fALFF in both DR and VTA, while a nominally significant positive correlation was found between EOT and fALFF in DR. However, none of these findings survived FDR correction (p_FDR_ > 0.05; ***Table 2***; for full results see ***Supplementary Table 2***).

**Table 2.** Associations between the neural activity amplitude measures and alexithymia scores.

| TAS Domain | Amplitude Measure | ROI | T-stat | p-value | FDR |
| --- | --- | --- | --- | --- | --- |
| DIF | fALFF | DR | -2.691 | 0.0094 ** | 0.0753 |
|  |  | VTA | -2.058 | 0.0443 * | 0.1688 |
| EOT | fALFF | DR | 2.083 | 0.0419 * | 0.2328 |

**TAS:** Toronto Alexithymia Scale; **DIF:** Difficulty Identifying Feelings; **EOT:** Externally Oriented Thinking; **(f)ALFF:** Fractional Amplitude of Low-Frequency Fluctuations; **ROI:** Region of Interest; **DR:** Dorsal Raphe; **VTA:** Ventral Tegmental Area
\* $p < 0.05$ ; \*\* $p < 0.01$

The majority of alexithymia scales, i.e. DIF, DDF and overall score, were however negatively associated with the functional connectivity between the ROIs and cortical regions (cluster-level FWE < 0.05; see ***Table 3***). In particular, negative associations were found for the functional connectivity of the DR and right LC with parietal regions involved in spatial cognition, reasoning, social cognition and working memory (Fan et al., 2016). Furthermore, similar alterations were observed for functional connections underlying social and semantic cognition, including the coupling of the right superior temporal gyrus with the left middle temporal gyrus (Wernicke’s area), and of the left LC with ipsilateral dorsomedial prefrontal cortex. Lastly, higher total alexithymia scores were further related to diminished functional connectivity between the right LC and the left ventromedial prefrontal cortex, an area associated with facial monitoring and discrimination (Fan et al., 2016). Conversely, two significant positive associations with EOT and DDF emerged, respectively, on the functional connectivity between VTA and the right inferior parietal lobule, and between DR and the left middle temporal gyrus (cluster-level FWE < 0.05). A visual representation of these findings is provided in ***Figure 2***. Although a nominally significant negative association between phonetic fluency and the mean functional connectivity of DR with the left postcentral gyrus and inferior parietal lobule was found (r = −0.263, p = 0.0423), it failed to survive the FDR < 0.05 correction (p_FDR_ = 0.2475) (see ***Supplementary Table 3***). As such, the observed results reflect to a great extent individual differences in emotional cognition, however, for certain connections, such as the coupling between the DR and parietal cortices, they may represent contributions from more generalised linguistic processing. The scatter plots with individual subject data are available in ***Supplementary Figure 2***.

**Figure 2.**
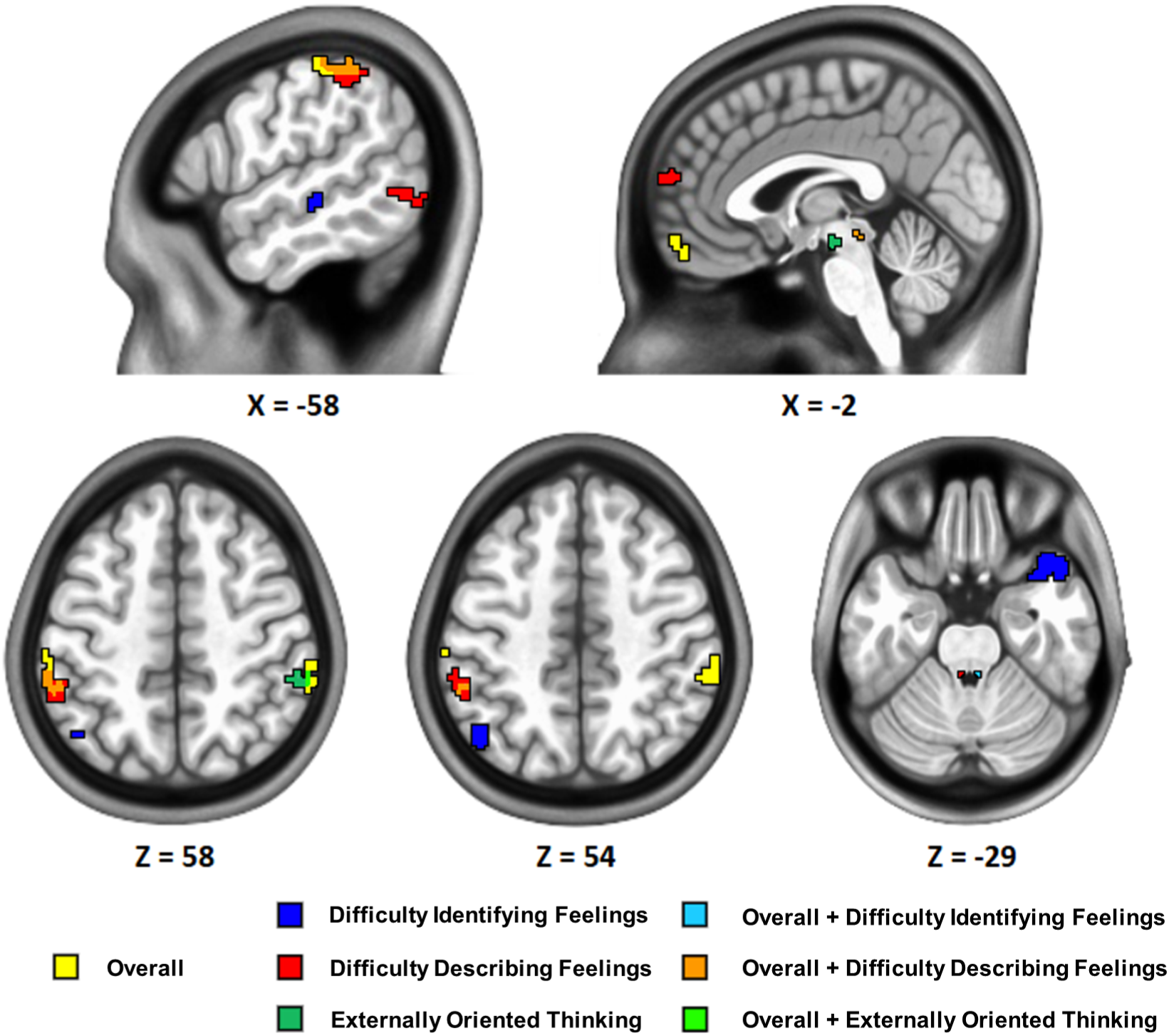
Visual representation of the significant resting-state functional connectivity findings color-coded by alexithymia scale.

**Table 3.** Significant associations between alexithymia scores and the resting-state functional connectivity patterns.

| TAS domain | ROI | MNI |  |  | Location | Voxels | F-stat | Pearson's R | Confidence Interval (95%) | Cohen's d |
| --- | --- | --- | --- | --- | --- | --- | --- | --- | --- | --- |
|  |  | x | y | z |  |  |  |  |  |  |
| DIF | R STG <sup>a</sup> | -64 | -25 | -5 | L middle temporal gyrus | 40 | 23.24 | 0.63 | 0.448 ; 0.762 | 1.623 |
|  | R LC | -46 | -66 | 46 | L inferior parietal lobule | 41 | 31.24 | -0.565 | -0.716 ; -0.363 | -1.370 |
| DDF | DR | -57 | -41 | 50 | L inferior parietal lobule | 59 | 24.44 | -0.583 | -0.729 ; -0.386 | -1.435 |
|  | DR | -60 | -61 | 0 | L middle temporal gyrus | 29 | 26.75 | -0.629 | -0.761 ; -0.446 | -1.618 |
|  | L LC | -5 | 60 | 18 | L dorsomedial prefrontal cortex | 37 | 28.49 | -0.638 | -0.768 ; -0.458 | -1.657 |
| EOT | VTA | 57 | -36 | 55 | R inferior parietal lobule | 33 | 32.42 | 0.56 | 0.357 ; 0.713 | 1.352 |
| Overall Score | DR | 62 | -34 | 50 | R inferior parietal lobule | 49 | 23.78 | -0.548 | -0.704 ; -0.341 | -1.310 |
|  | DR | -55 | -32 | 57 | L postcentral gyrus<br>L inferior parietal lobule | 48 | 26.51 | -0.633 | -0.764 ; -0.452 | -1.635 |
|  | R LC | -7 | 58 | -12 | B ventromedial prefrontal cortex | 35 | 31.48 | -0.603 | -0.743 ; -0.412 | -1.512 |

**TAS:** Toronto Alexithymia Scale; **DIF:** Difficulty Identifying Feelings; **DDF:** Difficulty Describing Feelings; **EOT:** Externally Oriented Thinking; **MNI:** Montreal Neurological Institute; **R:** Right; **L:** Left; **B:** Bilateral; **STG:** Superior Temporal Gyrus; **LC:** Locus Coeruleus; **DR:** Dorsal Raphe; **VTA:** Ventral Tegmental Area.
<sup>a</sup> This ROI pertains to the R STG0 subdivision as described in the work by Hoffman and Lambon-Ralph (2018).

### 3.2. Characteristics of specific anatomical connections

Direct anatomical pathways between areas highlighted in the functional connectivity analysis were found in at least one third of the sample for the following pairs of regions: DR and the left anterior inferior parietal lobule, DR and the right anterior inferior parietal lobule, and VTA and the right anterior inferior parietal lobule. In the case of the DR - left anterior inferior parietal lobule connection, a nominally significant positive association was found between the mean FA and DDF (t = 2.635, p = 0.0218), but it did not survive FDR correction (p_FDR_ = 0.0654). Additionally, a trend-level link emerged for the same connection between its functional connectivity and FA values (t = −1.8931, p = 0.0638). For a visualization of the specific tract please see ***Supplementary Figure 3***. No other findings reached nominal or corrected significance levels. Full results are available in ***Supplementary Tables 4 and 5***.

### 3.3. Graph theory-derived global and local network parameters

As shown in ***Figure 3***, network modularity was negatively associated with DIF, DDF and overall alexithymia score. On the same vein, the clustering coefficient of the left posterior middle temporal gyrus area (focal point in: x = −60, y = −61, z = 0) was negatively associated with DIF and overall alexithymia score. Neither parameter was correlated with the motion across the diffusion scan, nor with any of the semantic scores (p values > 0.05). No other results survived FDR < 0.05 correction (see ***Supplementary Table 6***).

**Figure 3.**
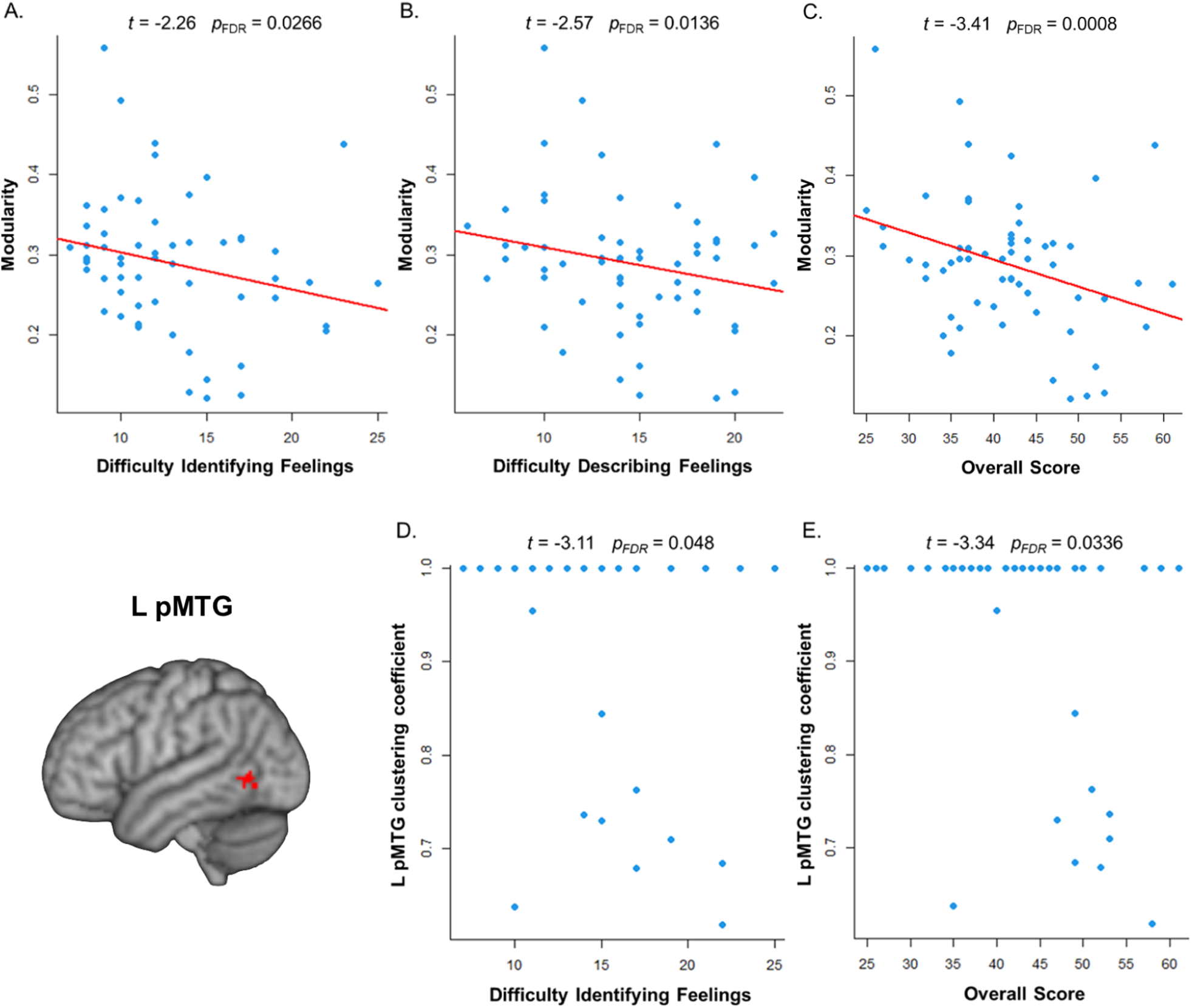
Significant associations of alexithymia scores with network modularity (A-C.) and the cluster coefficient of the left posterior middle temporal gyrus (D-E.; pMTG; MNI peak coordinates: x = −60, y = −61, z = 0).

### 3.4. Mediation results

Reappraisal data (ERQ; Abler & Kessler, 2009; Gross & John, 2003) was excluded from the mediation analysis as no relation was found between this emotion regulation strategy and the neuroimaging findings. On the other hand, as can be seen in ***Figure 4***, linear regression models determined significant positive associations for trait-anxiety (STAI) and the suppression strategy (ERQ) with DIF and DDF, respectively, and with alexithymia’s overall score. Subsequent linear regressions found significant associations for trait-anxiety and suppression with some of the functional connectivity findings. Trait-anxiety was positively related to the functional connectivity between the right superior temporal gyrus and the left middle temporal gyrus, and negatively associated with the functional connectivity between the right LC and left inferior parietal lobule. Additionally, habitual use of suppression strategy was linked with diminished functional connectivity between the left LC and the left dorsomedial prefrontal cortex (see ***Supplementary Tables 7 and 8***). Although significant effects were observed between the respective alexithymia measures and neuroimaging findings, no significant mediation effects of trait-anxiety nor suppression were found. Paired observations plots for the data used in the mediation models are available in ***Supplementary Figure 4***.

**Figure 4.**
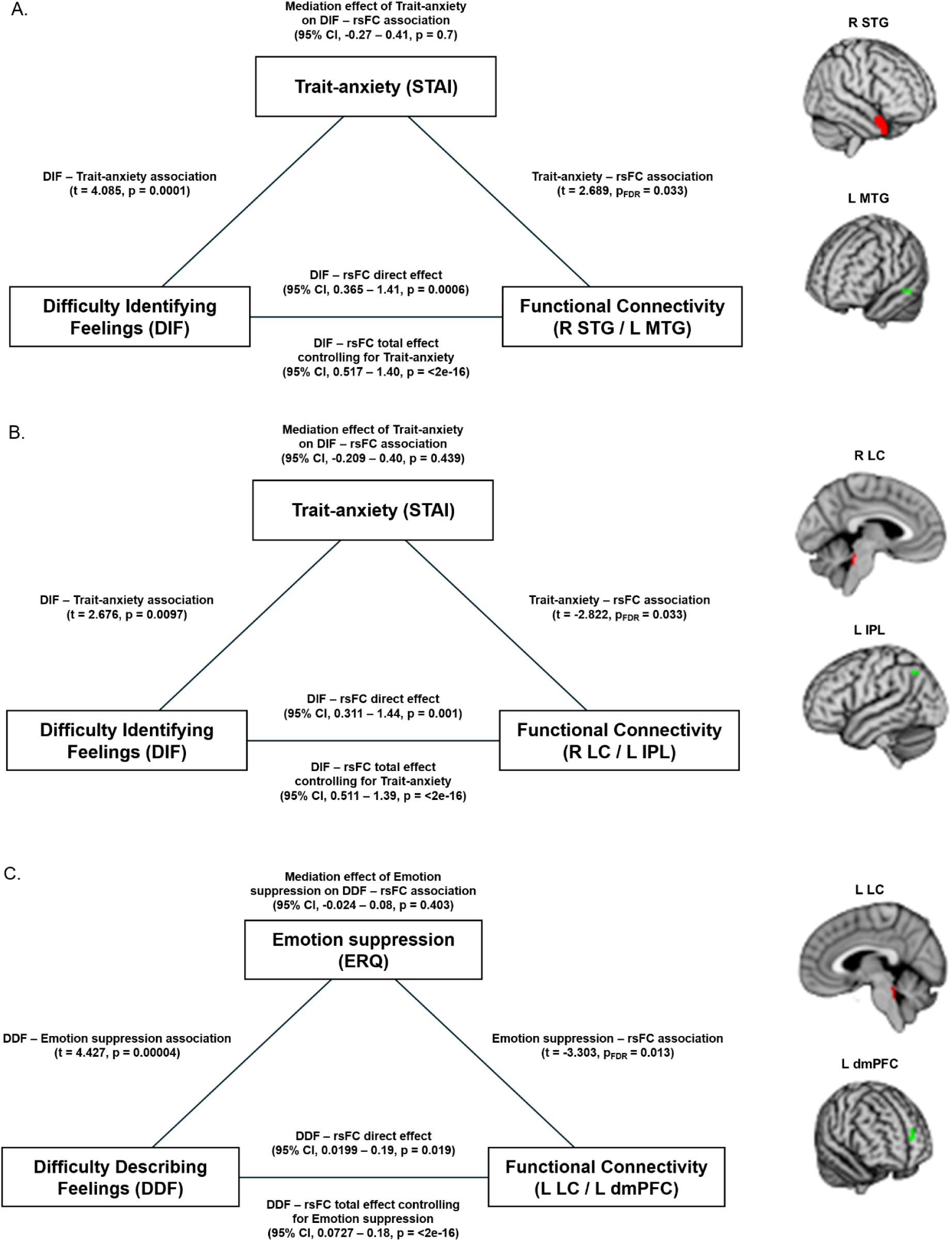
Mediation effects of trait-anxiety and habitual use of emotion suppression strategies on the association between alexithymia scales and resting-state functional connectivity (rsFC). **A.)** Trait-anxiety mediation plot for the rsFC between right superior temporal gyrus (R STG) and left middle temporal gyrus (L MTG). **B.)** Trait-anxiety mediation plot for the rsFC between the right locus coeruleus (R LC) and left inferior parietal lobule (L IPL). **C.)** Emotion suppression mediation plot for the rsFC between the left locus coeruleus (L LC) and the left dorsomedial prefrontal cortex (L dmPFC).

## 4. Discussion

In the current study, we aimed at increasing the understanding of alexithymia’s effects on the functional and structural organization patterns within the semantic and neuromodulatory brain networks underlying socio-emotional cognition. To this end, we employed the available rs-fMRI, dMRI and behavioural data of 60 healthy young adults from the *Max Planck Institute Mind-Brain-Body Dataset* (Babayan et al., 2019) to investigate the neural correlates of alexithymia in said networks.

Despite previous evidence pointing towards the existence of alterations in the semantic and neuromodulatory brain networks for alexithymia, as well as the relevant implications this bears for the accuracy in delineating the neural correlates of its highly comorbid conditions, such as anxiety and ASD, the matter at hand remains yet to be directly addressed in the literature. The current study aims to fill in this crucial gap of knowledge by demonstrating that the distinct dimensions of alexithymia (i.e. DIF, DDF and EOT) are all predominantly associated with unique patterns of functional coupling within the socio-emotional brain networks rather than localised resting-state activity. The importance of functional connectivity for understanding this phenomenon is further highlighted by the link between the dysconnectivity within the semantic system and DIF, underlining the peculiar contributions of semantic cognition to this process. On the structural level, these functional alterations are paralleled to a greater extent by the global and local organisation of the associated neural network, rather than the characteristics of specific anatomical connections. In particular, facets of alexithymia have been linked to lower network modularity and reduced local information processing around the left middle temporal gyrus. Importantly, the mediation analysis stresses that anxiety and emotion suppression do not act as a link between alexithymia and its neural correlates, emphasising the need to treat them as separate constructs.

In more detail, the functional connectivity analysis revealed positive and negative significant connections between the ROIs and multiple cortical areas for each alexithymia facet (see ***Table 3***). Greater difficulties identifying feelings were associated with lower connectivity between the right temporal pole (R STG0) and the left central middle temporal gyrus, a region underlying processes such as social cognition, semantics and auditory perception (Fan et al., 2016). This finding adds to the currently developing body of research that points towards a dysconnectivity pattern between the right temporal pole and socio-emotional processing network in alexithymia (Dobrushina et al., 2022). Interestingly, the connectivity between the DR and a more posterior area of the left middle temporal gyrus was positively correlated with difficulties describing feelings. Notably, a recent study on obsessive-compulsive disorder by Kim et al. (2019) found that higher functional connectivity between these areas correlated with unresponsiveness to selective serotonin reuptake inhibitors, and greater symptom severity, which has been associated in turn with greater difficulties describing feelings (Khosravani et al., 2017). Another finding relevant to this alexithymia facet was its negative association with the connectivity between the left LC and ipsilateral dorsomedial prefrontal cortex, a mentalizing-related region involved in social cognition and top-down emotion regulation (Eslinger et al., 2021; Fan et al., 2016, Ray & Zald, 2012). On a similar note, overall alexithymia decreased in relation to greater connectivity between the right LC and bilateral ventromedial prefrontal cortex, an area associated with facial monitoring and discrimination, and emotion regulation (Fan et al., 2016; Ray & Zald, 2012). Both these findings seem to support the idea that enhanced LC - prefrontal cortex connectivity supports the processing of emotional ambiguous cues, and is linked to better emotional health outcomes (Dave et al., 2024). Lastly, the results showed decreased functional connectivity between the neuromodulatory regions and bilateral inferior parietal regions involved in spatial and social cognition, reasoning and working memory (Fan et al., 2016) for all core aspects of alexithymia except externally oriented thinking, which was related to increased connectivity between VTA and the right inferior parietal lobule. While it has been proposed that the right inferior parietal lobule may facilitate self-directed thinking (Benedek et al., 2016), some studies suggest its general participation in switching between the internally- and externally-oriented attention (Uddin et al., 2019), with dopamine playing a crucial role in this process (Dang et al., 2012). However, relationships between dopamine and cognitive performance follow an inverted U-shaped profile (Dang et al., 2012; van Kempen et al., 2022). The current results, therefore, suggest that the functional pathway between VTA and the parietal cortex may preferentially respond to the processing of external stimuli in highly alexithymic individuals. In turn, the dysconnectivity of the inferior parietal lobe with DR and LC could possibly reflect diminished signalling of emotional cues to the attentional cortical areas, contributing to the observed deficits in their identification and naming. The notion that decreased serotonergic signalling may be linked to emotion processing, in specific the recognition of emotional states (Merens et al., 2007), is reinforced by the negative association found between alexithymia’s DIF domain and fALFF values in the DR, which, nevertheless, only reached trend-level significance.

As for the analysis of the structural connectivity between the pairs of regions from the functional analysis, it revealed a scarcity of direct anatomical connections, implicating that the functional coupling was more strongly mediated through indirect pathways (***Figure 1***). With the identified areas largely involved in complex, higher-order cognitions (Fan et al., 2016), the apparent negative results are in line with the reports of decreasing structure-function coupling from unimodal to transmodal cortices (Liu et al., 2022). Nevertheless, for the connection between DR and the left anterior inferior parietal lobule, which was identified in nearly all subjects, the current approach demonstrated trend-level associations of its mean FA with DDF, i.e. the same alexithymia dimension that was found in the functional analysis, and with the functional coupling itself.

The findings emerging from the graph analysis provided a further insight into the structural correlates of alexithymia within the identified network. We observed that greater modularity was associated with lower overall levels of alexithymia, and, particularly, with fewer difficulties identifying and describing feelings, indicating that the prevalence of these phenomena might be related to a loss of specialised neural processing, potentially reducing the network’s ability to distinguish between different types of information. This characteristic is strikingly reminiscent of a key aspect of alexithymia, i.e. not being able to differentiate between emotions and bodily sensations (Brewer et al., 2016). Prior literature (Guassi Moreira et al., 2021; Richardson et al., 2018) has linked greater modularity in the default mode network, which plays a key role in social and self-referential cognition, with higher emotion regulation and mentalizing abilities. On the same note, a study by Liemburg et al. (2012) found diminished connectivity within this network, accompanied by stronger functional connectivity with sensory and prefrontal areas in alexithymic individuals. In addition to the associations with the global network parameters, we observed that the clustering coefficient of the left posterior middle temporal gyrus was similarly negatively related to the overall alexithymia scores and DIF. This finding complements the associations with DDF observed for this semantic node in the functional analysis. Combined, these results hint at the crucial role of the deficits in emotional semantic processing in alexithymia, which might be, in fact, the primary mechanism preventing such individuals from effective emotion regulation (Lieberman et al., 2007).

Importantly for these considerations, the mediation analysis indicated that trait-anxiety and habitual use of emotion suppression strategies did not act as a link between alexithymia and associated neural correlates, providing further evidence for the conceptualization of alexithymia as an independent construct (Bernhardt et al., 2014; Kinnaird et al., 2019; Marchesi et al., 2000). However, as the current work has been performed on a subclinical sample, these findings cannot be generalised to patient populations. Potential future avenues of research could focus on studying the predictive value of the observed neural patterns associated with alexithymia in regards to the future development of anxiety and mood symptomatology on a similar subclinical sample. Another topic of interest that future studies should examine is the possibility that alexithymia may differentially affect clinical and subclinical populations. Furthermore, these studies could benefit from including a bigger sample size, as we may have overlooked potential results due to a lack of statistical power. Lastly, an inherent characteristic of the diffusion MRI-based tractography employed in this study is that the reconstruction of longer streamlines, including interhemispheric connections, often lacks accuracy (Yeh et al., 2021).

## 5. Conclusions

In conclusion, the findings emerging from this study support the notion of alexithymia as an independent construct with unique functional and structural neural correlates. Furthermore, we provide evidence of distinct functional mapping patterns between alexithymia’s domains in the social semantics network and the neuromodulatory circuitry that could potentially explain its symptomatology. Lastly, we suggest that the characteristics of the observed network constitute the mechanism preventing alexithymic individuals from effectively regulating their emotions.

## Data availability

ROIs for the functional analysis and unthresholded statistical maps generated for this study are available on the following NeuroVault repository: https://identifiers.org/neurovault.collection:22253. The original data used for the analysis can be found in the associated data paper (Babayan et al., 2019). The scripts used for processing the functional data after preprocessing with fMRIprep (Esteban et al., 2019), together with the scripts used in the diffusion analysis are available at the following GitHub repository: https://github.com/Montolim/10.1101-2025.05.23.655721.

## CRediT author statement

MIM: Conceptualization, Formal analysis, Writing - Original Draft, Visualization. MV: Conceptualization, Writing - Review & Editing, Supervision, Funding acquisition. MRZ: Conceptualization, Formal analysis, Writing - Review & Editing, Supervision.

## Ethical statement

The study was performed using an openly available, fully anonymised dataset (Babayan et al., 2019), and as such was not subjected to the ethical committee approval.

