## Supplementary Material for "Socio-emotional difficulties observed in alexithymia reflect altered interactions of the semantic and monoaminergic neuromodulatory brain networks"

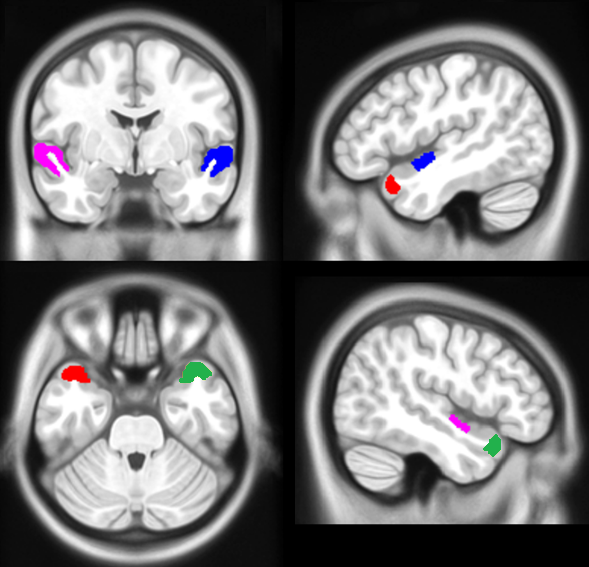

**Supplementary Figure 1. Representation of the STG subdivisions employed in our study as described in the work by Hoffman and Lambon-Ralph (2018).** Red: left STG0; Blue: left STG2; Green: right STG0; Purple: right STG2. Coordinates for the slices: left (-47, -3, -29); right (47, -3, -29) .

**Supplementary Figure 2. Scatter plots for the significant associations between resting-state functional connectivity values and Difficulty Describing Feelings (A-C.), Difficulty Identifying Feelings (D-E.), Externally Oriented Thinking (F.), and Overall alexithymia scores (G-I.).**

**L:** left; **R:** right; **B:** bilateral; **DR:** Dorsal Raphe; **IPL:** Inferior Parietal Lobule; **MTG:** Middle Temporal Gyrus; **LC:** Locus Coeruleus; **DMPFC:** Dorsomedial Prefrontal Cortex; **STG:** Superior Temporal Gyrus; **VTA:** Ventral Tegmental Area; **PG:** Postcentral Gyrus; **VMPFC:** Ventromedial Prefrontal Cortex.

**R STG0** refers to the superior temporal gyrus subdivision described in the work by Hoffman and Lambon-Ralph (2018).

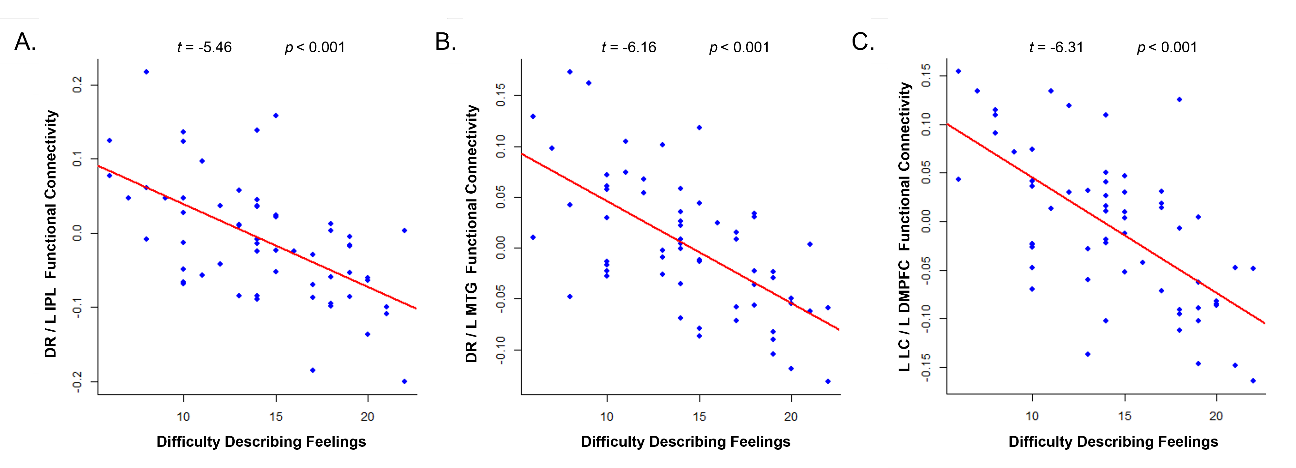

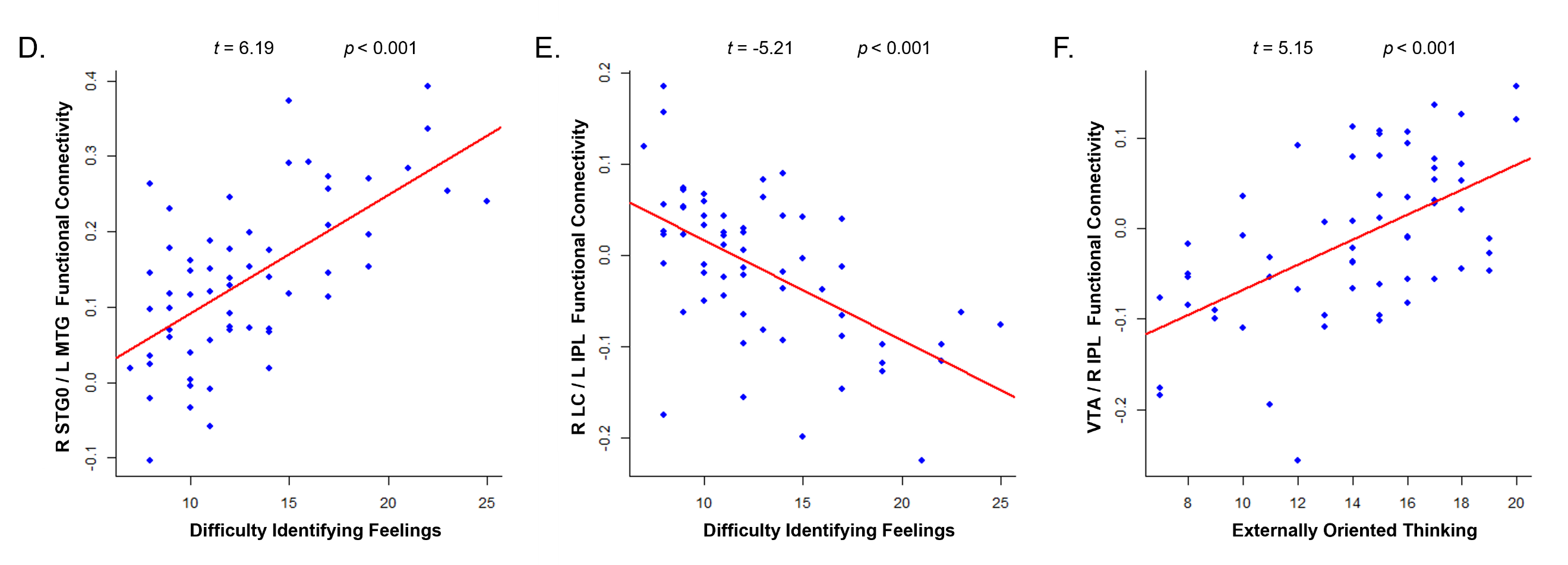

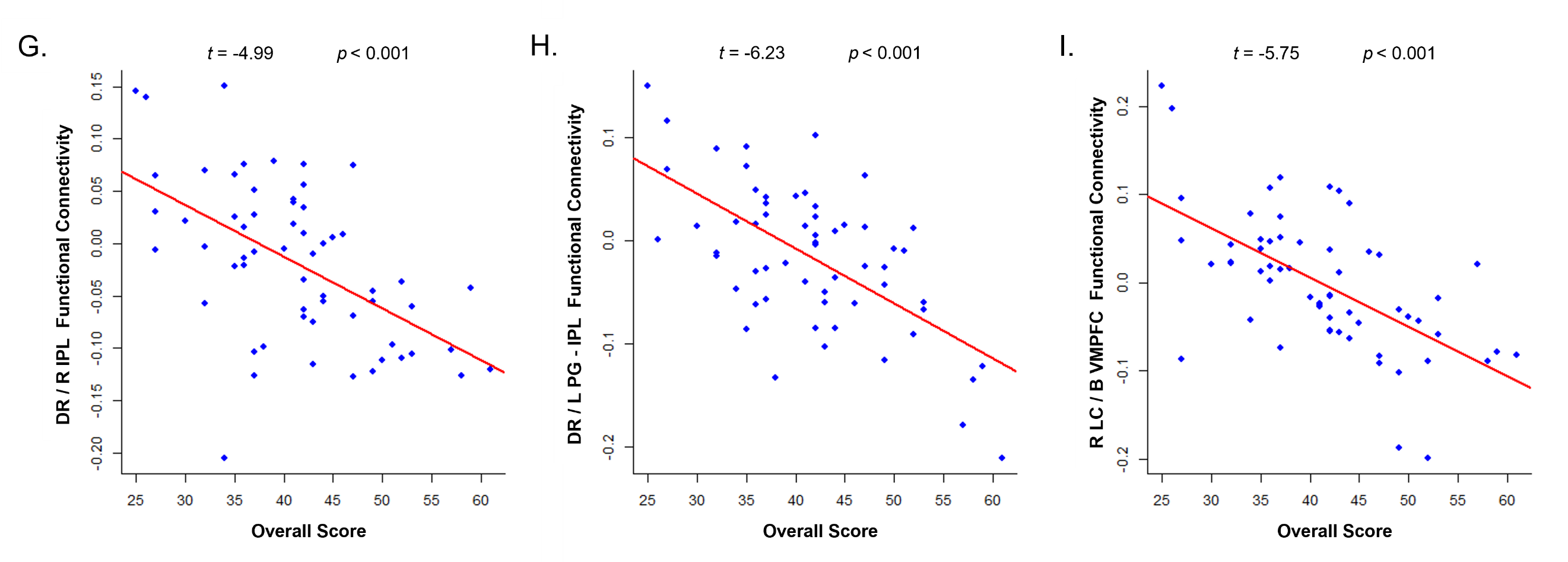

**Supplementary Figure 3. Visualization of the tracts linking the Dorsal Raphe with the left anterior Inferior Parietal Lobule (L aIPL) for two representative subjects.** Center of mass coordinates, MNI: **L aIPL** (-56, -38, 51). **L aIPL** includes two overlapping areas from the functional analysis: **L IPL / PCG** (Postcentral Gyrus) (peak activity coordinates: -55, -32, 57) and **L IPL** (peak activity coordinates: -57, -41, 50).

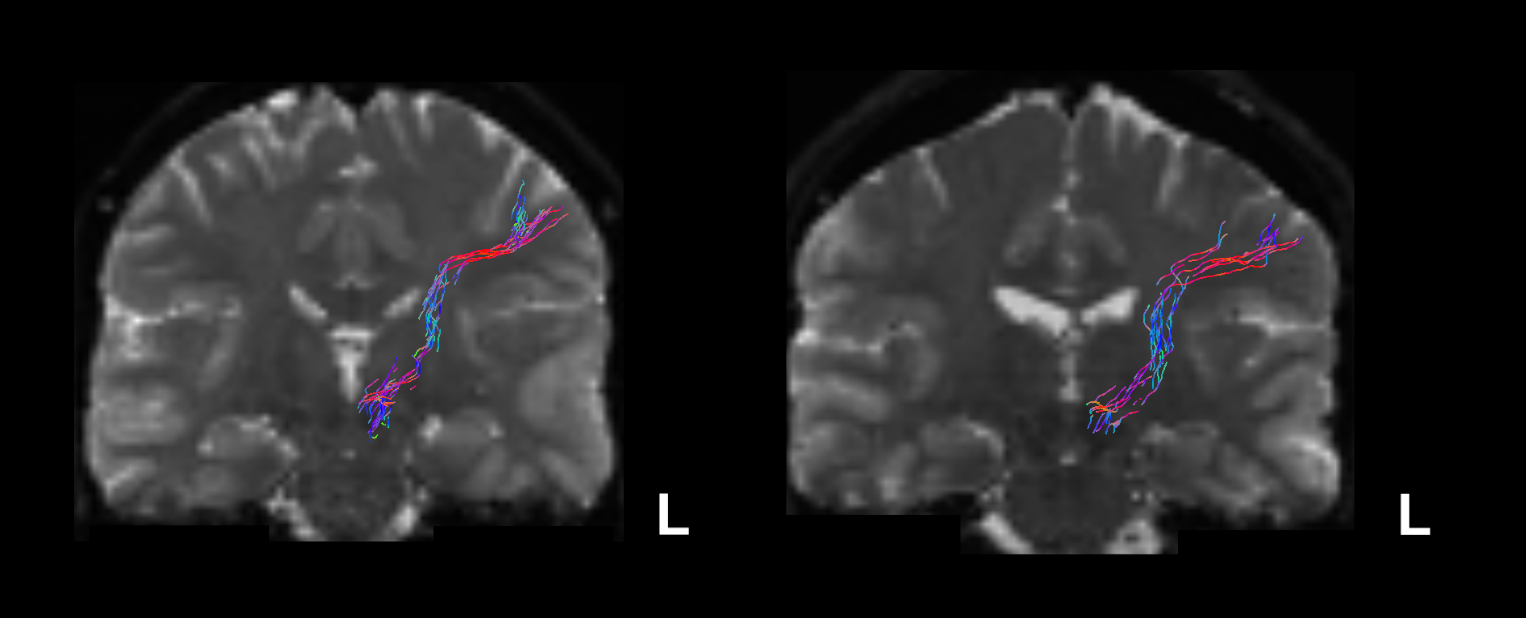

**Subject - 010163**

**Subject - 010015**

**Supplementary Figure 4. Standardised (z-score) paired observations plots for the mediation models. L:** left; **R:** right; **STG:** Superior Temporal Gyrus; **MTG:** Middle Temporal Gyrus; **LC:** Locus Coeruleus; **IPL:** Inferior Parietal Lobule; **DMPFC:** Dorsomedial Prefrontal Cortex.

**R STG0** refers to the superior temporal gyrus subdivision described in the work by Hoffman and Lambon-Ralph (2018).

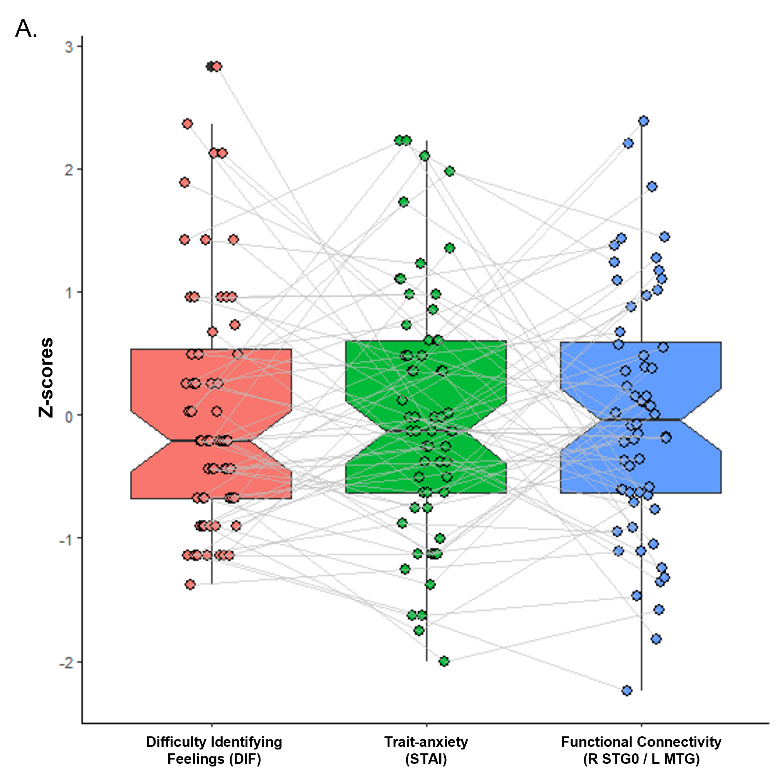

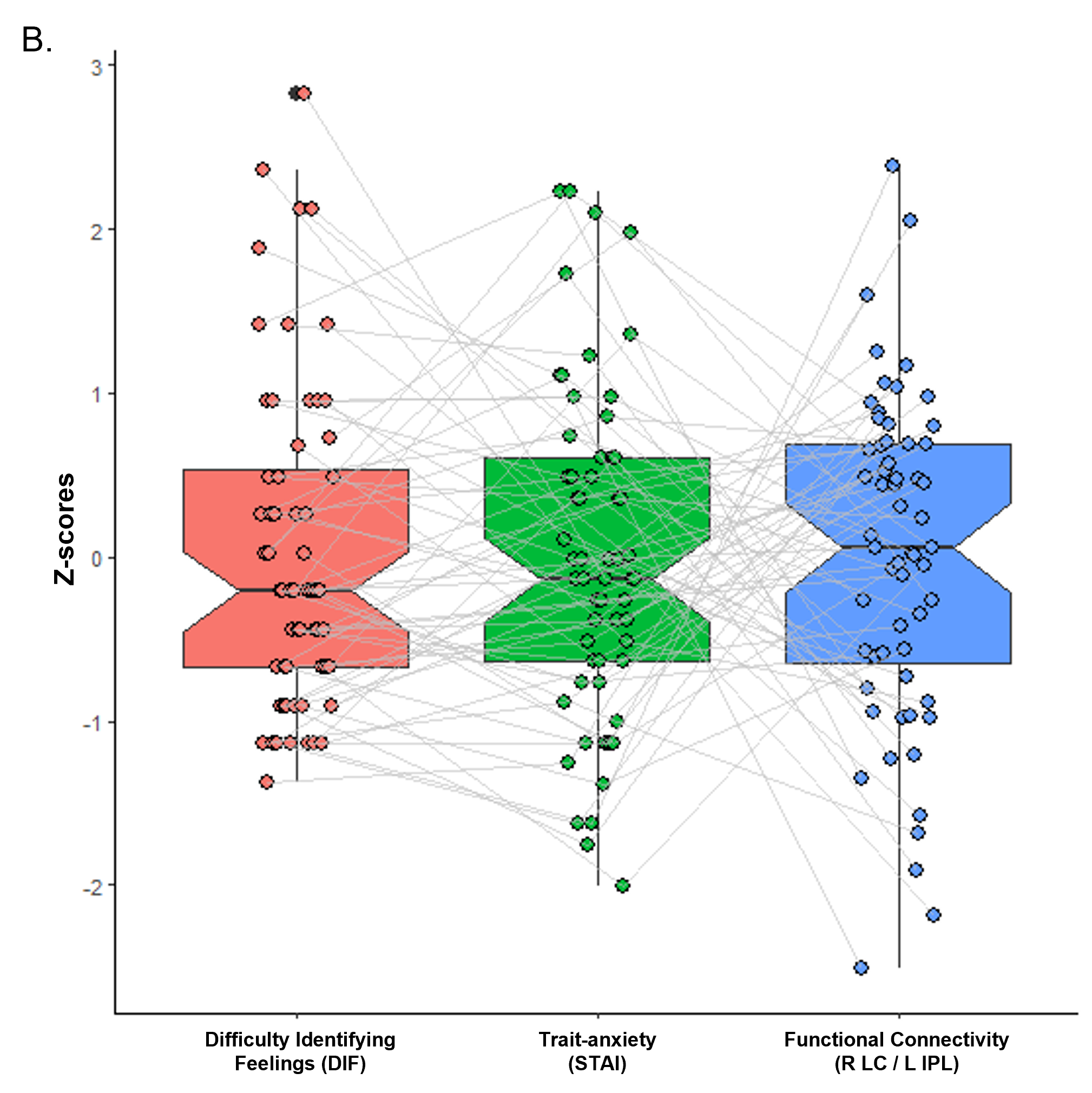

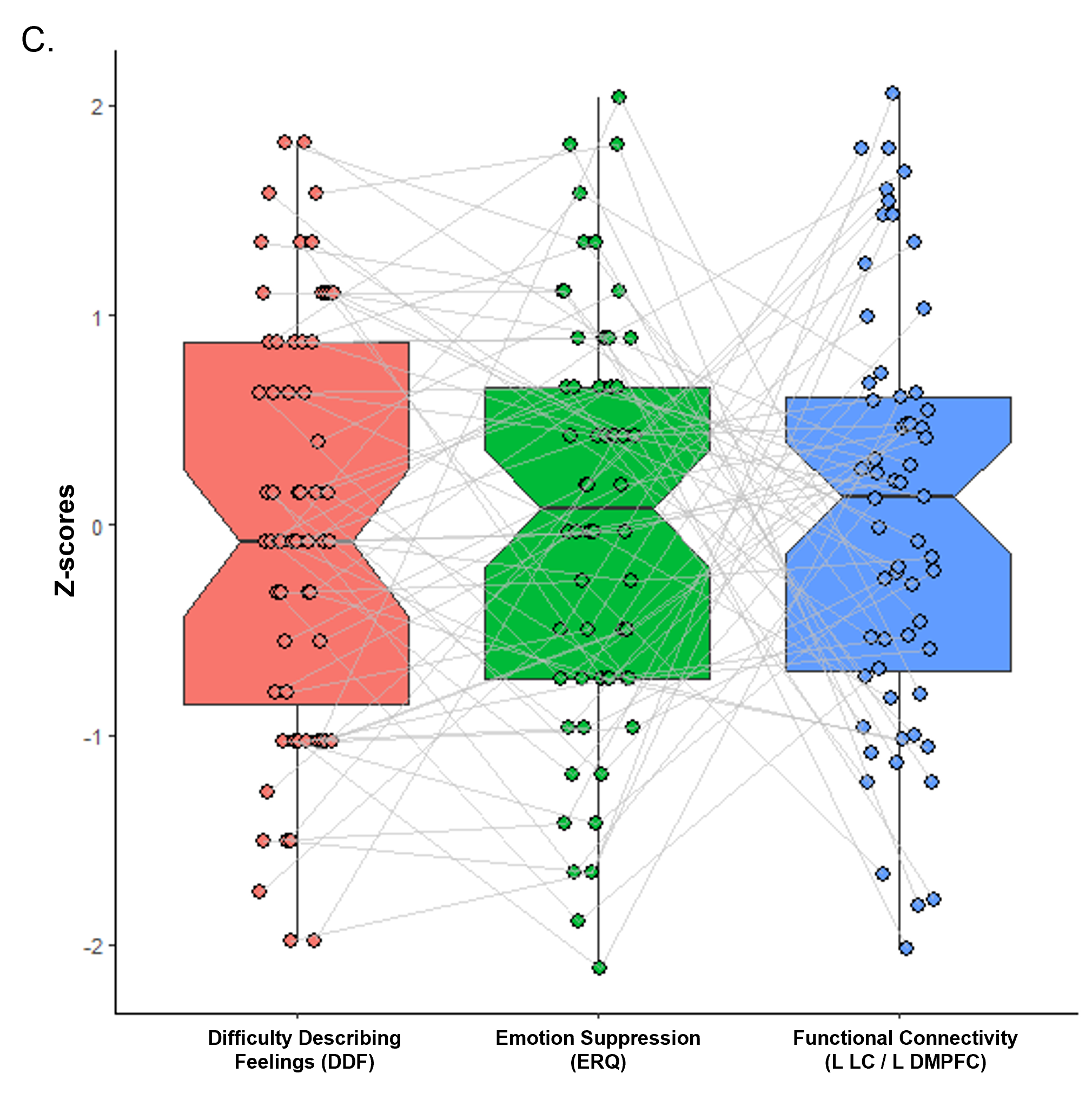

**Supplementary Table 1.**

Graph theory measures extracted from the structural network consisting of regions functionally associated with alexithymia in the current study.

| **Measure** | **Definition** |
| --- | --- |
| Weighted clustering coefficient | An extension of unweighted clustering coefficient, i.e. the proportion of the number of links between the node and its immediate neighbours divided by the number of links that could possibly exist between them, that takes into account the edge weights (Barrat et al., 2004). |
| Betweenness centrality | Number of the shortest paths in the graph that pass through the node (Freeman, 1979). |
| Global efficiency | Average of the inverse of the distances between all pairs of nodes (Latora and Marchiori, 2001) |
| Modularity | Extent to which a graph can be divided into clearly separated communities (Traag et al., 2019). |

**Supplementary Table 2.**

Associations between amplitude measures and alexithymia scores for each ROI.

| **TAS Domain** | **Amplitude Measure** | **ROI** | **T-stat** | **p-value** | **FDR** |
| --- | --- | --- | --- | --- | --- |
| DIF | ALFF | DR | 0.350 | 0.7278 | 0.7278 |
|  |  | VTA | 0.734 | 0.4663 | 0.6286 |
|  |  | L LC | 0.671 | 0.5050 | 0.6286 |
|  |  | R LC | 0.601 | 0.5500 | 0.6286 |
|  |  | R STG0 ^a^ | -0.958 | 0.3422 | 0.6286 |
|  |  | R STG2 ^b^ | -1.207 | 0.2326 | 0.6286 |
|  |  | L STG0 ^a^ | -0.708 | 0.4820 | 0.6286 |
|  |  | L STG2 ^b^ | -0.789 | 0.4340 | 0.6286 |
|  | fALFF | DR | -2.691 | 0.0094** | 0.0753 |
|  |  | VTA | -2.058 | 0.0443* | 0.1688 |
|  |  | L LC | -0.238 | 0.8130 | 0.8130 |
|  |  | R LC | -0.288 | 0.7747 | 0.8130 |
|  |  | R STG0 ^a^ | -1.235 | 0.2220 | 0.3552 |
|  |  | R STG2 ^b^ | -1.331 | 0.1886 | 0.3552 |
|  |  | L STG0 ^a^ | -1.895 | 0.0633 | 0.1688 |
|  |  | L STG2 ^b^ | -0.951 | 0.3458 | 0.4611 |
| DDF | ALFF | DR | 0.428 | 0.6704 | 0.7662 |
|  |  | VTA | 1.379 | 0.1734 | 0.6231 |
|  |  | L LC | 0.802 | 0.4260 | 0.6231 |
|  |  | R LC | 0.736 | 0.4650 | 0.6231 |
|  |  | R STG0 ^a^ | -0.732 | 0.4673 | 0.6231 |
|  |  | R STG2 ^b^ | -1.283 | 0.2050 | 0.6231 |
|  |  | L STG0 ^a^ | -0.745 | 0.4590 | 0.6231 |
|  |  | L STG2 ^b^ | 0.146 | 0.8840 | 0.8840 |
|  | fALFF | DR | -0.720 | 0.4750 | 0.5793 |
|  |  | VTA | -0.849 | 0.4000 | 0.5793 |
|  |  | L LC | -1.111 | 0.2710 | 0.5793 |
|  |  | R LC | -0.784 | 0.4360 | 0.5793 |
|  |  | R STG0 ^a^ | -0.168 | 0.8670 | 0.8670 |
|  |  | R STG2 ^b^ | -1.273 | 0.2083 | 0.5793 |
|  |  | L STG0 ^a^ | -1.289 | 0.2030 | 0.5793 |
|  |  | L STG2 ^b^ | -0.668 | 0.5069 | 0.5793 |
| EOT | ALFF | DR | 0.226 | 0.8221 | 0.8221 |
|  |  | VTA | -0.363 | 0.7177 | 0.8202 |
|  |  | L LC | 0.382 | 0.7040 | 0.8202 |
|  |  | R LC | 1.248 | 0.2170 | 0.4597 |
|  |  | R STG0 ^a^ | 1.074 | 0.2873 | 0.4597 |
|  |  | R STG2 ^b^ | 1.151 | 0.2546 | 0.4597 |
|  |  | L STG0 ^a^ | 1.212 | 0.2310 | 0.4597 |
|  |  | L STG2 ^b^ | 1.799 | 0.0776 | 0.4597 |
|  | fALFF | DR | 2.083 | 0.0419* | 0.2328 |
| EOT | fALFF | VTA | 1.176 | 0.2450 | 0.3267 |
|  |  | L LC | 0.970 | 0.3360 | 0.3840 |
|  |  | R LC | 0.546 | 0.5873 | 0.5873 |
|  |  | R STG0 ^a^ | 1.465 | 0.1490 | 0.2980 |
|  |  | R STG2 ^b^ | 1.250 | 0.2166 | 0.3267 |
|  |  | L STG0 ^a^ | 1.741 | 0.0873 | 0.2328 |
|  |  | L STG2 ^b^ | 1.854 | 0.0691 | 0.2328 |
| Overall Score | ALFF | DR | 0.490 | 0.6262 | 0.7957 |
|  |  | VTA | 0.898 | 0.3730 | 0.7957 |
|  |  | L LC | 0.910 | 0.3667 | 0.7957 |
|  |  | R LC | 1.217 | 0.2290 | 0.7957 |
|  |  | R STG0 ^a^ | -0.392 | 0.6962 | 0.7957 |
|  |  | R STG2 ^b^ | -0.753 | 0.4545 | 0.7957 |
|  |  | L STG0 ^a^ | -0.216 | 0.8300 | 0.8300 |
|  |  | L STG2 ^b^ | 0.423 | 0.6740 | 0.7957 |
|  | fALFF | DR | -0.797 | 0.4290 | 0.8898 |
|  |  | VTA | -0.945 | 0.3490 | 0.8898 |
|  |  | L LC | -0.253 | 0.8010 | 0.9681 |
|  |  | R LC | -0.300 | 0.7650 | 0.9681 |
|  |  | R STG0 ^a^ | -0.091 | 0.9280 | 0.9681 |
|  |  | R STG2 ^b^ | -0.769 | 0.4449 | 0.8898 |
|  |  | L STG0 ^a^ | -0.850 | 0.3990 | 0.8898 |
|  |  | L STG2 ^b^ | -0.040 | 0.9681 | 0.9681 |
| **TAS:** Toronto Alexithymia Scale; **DIF:** Difficulty Identifying Feelings; **DDF:** Difficulty Describing Feelings; **EOT:** Externally Oriented Thinking; **ALFF:** Amplitude of Low-Frequency Fluctuations; **fALFF:** fractional ALFF; **ROI:** Region of Interest; **L:** Left; **R:** Right; **DR:** Dorsal Raphe; **VTA:** Ventral Tegmental Area; **LC:** Locus Coeruleus; **STG:** Superior Temporal Gyrus; **FDR:** False-Discovery Rate.  * p < 0.05; ** p < 0.01  ^a^ This ROI pertains to the STG0 subdivision as described in the work by Hoffman and Lambon-Ralph (2018).  ^b^ This ROI pertains to the STG2 subdivision as described in the work by Hoffman and Lambon-Ralph (2018). | | | | | |

**Supplementary Table 3.**

Associations between identified resting-state functional connectivity patterns and general semantic cognition.

| **Connection** | **RWT_1 (r)** | **RWT_1 (p)** | **RWT_1 (FDR)** | **RWT_13 (r)** | **RWT_13 (p)** | **RWT_13 (FDR)** | **WST_1 (r)** | **WST_1 (p)** | **WST_1 (FDR)** |
| --- | --- | --- | --- | --- | --- | --- | --- | --- | --- |
| DR - L IPL | -0.183 | 0.16165 | 0.29097 | -0.0718 | 0.589835 | 0.8154079 | -0.0064 | 0.96371 | 0.96371 |
| DR - L MTG | -0.1024 | 0.438048 | 0.5866881 | -0.0471 | 0.721392 | 0.8154079 | -0.0713 | 0.589835 | 0.8847525 |
| L LC - L dmPFC | -0.2329 | 0.074475 | 0.24753 | 0.1107 | 0.39977 | 0.8154079 | 0.0242 | 0.854377 | 0.96371 |
| R STG ^a^ - L MTG | 0.1879 | 0.150522 | 0.29097 | 0.1688 | 0.197298 | 0.8154079 | 0.2441 | 0.060168 | 0.541512 |
| R LC - L IPL | 0.0234 | 0.859142 | 0.859142 | 0.0464 | 0.724807 | 0.8154079 | -0.1077 | 0.415803 | 0.8847525 |
| VTA - R IPL | -0.0985 | 0.456313 | 0.5866881 | -0.0581 | 0.659802 | 0.8154079 | -0.1052 | 0.424622 | 0.8847525 |
| DR - R IPL | -0.2269 | 0.08251 | 0.24753 | -0.1963 | 0.133389 | 0.8154079 | -0.0734 | 0.579373 | 0.8847525 |
| DR - L IPL / PCG | -0.263 | 0.042333* | 0.24753 | -0.1347 | 0.307379 | 0.8154079 | 0.0182 | 0.890223 | 0.96371 |
| R LC - L vmPFC | 0.0311 | 0.813519 | 0.859142 | -0.0297 | 0.825907 | 0.825907 | -0.0799 | 0.548508 | 0.8847525 |
| **L:** Left; **R:** Right; **DR:** Dorsal Raphe; **LC:** Locus Coeruleus; **STG:** Superior Temporal Gyrus; **VTA:** Ventral Tegmental Area; **IPL:** Inferior Parietal Lobule; **MTG:** Middle Temporal Gyrus; **dmPFC:** dorsomedial Prefrontal Cortex; **PCG:** Postcentral Gyrus; **vmPFC:** ventromedial Prefrontal Cortex; **RWT:** Regensburger Wortflüssigkeits-Test; **WST:** Wortschatztest; **FDR:** False-Discovery Rate.  * p < 0.05  ^a^ This ROI pertains to the R STG0 subdivision as described in the work by Hoffman and Lambon-Ralph (2018). | | | | | | | | | |

**Supplementary Table 4.**

Associations between structural measures and alexithymia for the functional connections with a structural correlate present in more than one third of the sample.

| **TAS Domain** | **Connection** | **Structural Measure** | **T-stat** | **p-value** | **FDR** |
| --- | --- | --- | --- | --- | --- |
| DIF | DR – L aIPL  (-56, -38, 51) | FA | 0.193 | 0.8480 | 0.8480 |
|  |  | ADC | 0.958 | 0.3430 | 0.8480 |
|  |  | streamlines | -0.407 | 0.6855 | 0.8480 |
|  | DR – R aIPL | FA | 0.312 | 0.7570 | 0.7570 |
|  |  | ADC | -0.348 | 0.7310 | 0.7570 |
|  |  | streamlines | -0.625 | 0.5370 | 0.7570 |
|  | VTA – R aIPL | FA | 0.358 | 0.7247 | 0.7247 |
|  |  | ADC | 0.472 | 0.6420 | 0.7247 |
|  |  | streamlines | 0.424 | 0.6770 | 0.7247 |
| DDF | DR – L aIPL  (-56, -38, 51) | FA | 2.365 | 0.0218* | 0.0654 |
|  |  | ADC | -1.895 | 0.0637 | 0.0956 |
|  |  | streamlines | 0.077 | 0.9386 | 0.9386 |
|  | DR – R aIPL | FA | 1.095 | 0.2830 | 0.4425 |
|  |  | ADC | -0.772 | 0.4460 | 0.4460 |
|  |  | streamlines | 1.066 | 0.2950 | 0.4425 |
|  | VTA – R aIPL | FA | 1.067 | 0.3000 | 0.3000 |
|  |  | ADC | -1.560 | 0.1360 | 0.2040 |
|  |  | streamlines | 1.561 | 0.1359 | 0.2040 |
| EOT | DR – L aIPL  (-56, -38, 51) | FA | 0.778 | 0.4400 | 0.7680 |
|  |  | ADC | 0.296 | 0.7680 | 0.7680 |
|  |  | streamlines | 0.360 | 0.7201 | 0.7680 |
|  | DR – R aIPL | FA | -0.595 | 0.5570 | 0.5570 |
|  |  | ADC | -0.986 | 0.3320 | 0.5570 |
|  |  | streamlines | -0.808 | 0.4258 | 0.5570 |
|  | VTA – R aIPL | FA | -0.499 | 0.6236 | 0.7640 |
|  |  | ADC | -0.365 | 0.7190 | 0.7640 |
|  |  | streamlines | 0.305 | 0.7640 | 0.7640 |
| Overall Score | DR – L aIPL  (-56, -38, 51) | FA | 1.596 | 0.1170 | 0.3510 |
|  |  | ADC | -0.275 | 0.7840 | 0.9790 |
|  |  | streamlines | -0.026 | 0.9790 | 0.9790 |
|  | DR – R aIPL | FA | 0.441 | 0.6630 | 0.8460 |
|  |  | ADC | -1.088 | 0.2860 | 0.8460 |
|  |  | streamlines | -0.196 | 0.8460 | 0.8460 |
|  | VTA – R aIPL | FA | 0.554 | 0.5860 | 0.5860 |
|  |  | ADC | -0.829 | 0.4180 | 0.5860 |
|  |  | streamlines | 1.380 | 0.1845 | 0.5535 |
| **TAS:** Toronto Alexithymia Scale; **DIF:** Difficulty Identifying Feelings; **DDF:** Difficulty Describing Feelings; **EOT:** Externally Oriented Thinking; **L:** Left; **R:** Right; **DR:** Dorsal Raphe; **VTA:** Ventral Tegmental Area; **aIPL:** anterior Inferior Parietal Lobule; **FA:** Fractional Anisotropy; **ADC:** Apparent Diffusion Coefficient; **FDR:** False-Discovery Rate.  * p < 0.05 | | | | | |

**Supplementary Table 5.**

Associations between structural measures and functional connectivity values for the functional connections with a structural correlate present in more than one third of the sample.

| **Connection** | **Structural Measure** | **T-stat** | **p-value** |
| --- | --- | --- | --- |
| DR – L aIPL  (-56, -38, 51) | FA | -1.893 | 0.0638 |
|  | ADC | 0.144 | 0.8861 |
|  | streamlines | 0.624 | 0.5352 |
| DR – R aIPL | FA | 0.145 | 0.8858 |
|  | ADC | 0.579 | 0.5670 |
|  | streamlines | 0.147 | 0.8845 |
| VTA – R aIPL | FA | 0.917 | 0.3712 |
|  | ADC | 0.786 | 0.4421 |
|  | streamlines | 1.028 | 0.3177 |
| **L**: Left; R: Right; **DR**: Dorsal Raphe; **VTA**: Ventral Tegmental Area; **aIPL**: anterior Inferior Parietal Lobule; **FA**: Fractional Anisotropy; **ADC**: Apparent Diffusion Coefficient. | | | |

**Supplementary Table 6.**

Associations between the graph measures and alexithymia domains

| **Graph Measure** | | **TAS DIF** | | | **TAS DDF** | | | **TAS EOT** | | | **TAS Overall Score** | | |
| --- | --- | --- | --- | --- | --- | --- | --- | --- | --- | --- | --- | --- | --- |
|  |  | **t** | **p** | **FDR** | **t** | **p** | **FDR** | **t** | **p** | **FDR** | **t** | **p** | **FDR** |
| Mean Cluster Coefficient | | 1.06 | 0.2828 | 0.2828 | 0.21 | 0.8336 | 0.8336 | 0.73 | 0.4570 | 0.4570 | 0.98 | 0.3262 | 0.3262 |
| Global Efficiency | | 0 | 0.9956 | 0.9956 | 1.69 | 0.0986 | 0.0986 | 0.36 | 0.7106 | 0.7106 | 0.98 | 0.3266 | 0.3266 |
| Modularity | | -2.26 | **0.0266*** | **0.0266*** | -2.57 | **0.0136*** | **0.0136*** | -1.8 | 0.0804 | 0.0804 | -3.41 | **0.0008***** | **0.0008***** |
| Clustering Coefficient | B vmPFC | 1.29 | 0.2090 | 0.5650 | 0.95 | 0.3476 | 0.6952 | -0.28 | 0.7840 | 0.7924 | 1.03 | 0.2904 | 0.6970 |
|  | DR | 0.47 | 0.6448 | 0.8234 | -0.15 | 0.8852 | 0.8992 | -1.14 | 0.2586 | 0.5515 | -0.31 | 0.7522 | 0.9220 |
|  | L dmPFC | 1.47 | 0.1398 | 0.5592 | 1.36 | 0.1686 | 0.5058 | 0.92 | 0.3590 | 0.5515 | 1.88 | 0.0638 | 0.3828 |
|  | L aIPL ^b^ | 2.18 | 0.0324* | 0.1944 | 1.96 | 0.0546 | 0.2880 | -0.83 | 0.4136 | 0.5515 | 1.72 | 0.0980 | 0.3920 |
|  | L IPL ^a^ | -0.5 | 0.6154 | 0.8234 | -0.62 | 0.5546 | 0.8319 | 1.1 | 0.2776 | 0.5515 | -0.1 | 0.9220 | 0.9220 |
|  | L LC | 0.04 | 0.9648 | 0.9648 | 0.12 | 0.8992 | 0.8992 | -1.78 | 0.0830 | 0.498 | -0.67 | 0.4908 | 0.9220 |
|  | L cMTG ^a^ | 0.6 | 0.5526 | 0.8234 | 0.83 | 0.4104 | 0.7035 | -1.31 | 0.1970 | 0.5515 | 0.17 | 0.8654 | 0.9220 |
|  | L pMTG ^a^ | -3.11 | **0.0040**** | **0.0480*** | -2.39 | 0.0208* | 0.2496 | -0.94 | 0.3452 | 0.5515 | -3.34 | **0.0028**** | **0.0336*** |
|  | R aIPL | -0.55 | 0.5882 | 0.8234 | -0.2 | 0.8412 | 0.8992 | 1.88 | 0.0702 | 0.498 | 0.4 | 0.6854 | 0.9220 |
|  | R LC | -0.32 | 0.7548 | 0.8234 | -0.18 | 0.8618 | 0.8992 | 0.27 | 0.7924 | 0.7924 | -0.14 | 0.8904 | 0.9220 |
|  | R STG ^c^ | 1.22 | 0.2354 | 0.56496 | -1.18 | 0.2496 | 0.5990 | 0.38 | 0.7062 | 0.7924 | 0.22 | 0.8264 | 0.9220 |
|  | VTA | -0.39 | 0.7070 | 0.8234 | -1.83 | 0.0720 | 0.288 | -0.83 | 0.4128 | 0.5515 | -1.46 | 0.1444 | 0.4332 |
| Betweenness Centrality | B vmPFC | 0.47 | 0.6430 | 0.7834 | 0.21 | 0.8378 | 0.8378 | 0.05 | 0.9564 | 0.9564 | 0.38 | 0.7114 | 0.7761 |
|  | DR | -0.09 | 0.9282 | 0.9282 | 0.35 | 0.7212 | 0.7876 | 2.14 | 0.0384* | 0.4608 | 1.02 | 0.3008 | 0.5133 |
|  | L dmPFC | -0.47 | 0.6528 | 0.7834 | 0.9 | 0.3688 | 0.6322 | 0.71 | 0.4768 | 0.8174 | 0.5 | 0.6098 | 0.7318 |
|  | L aIPL ^b^ | -2.75 | 0.0082** | 0.0624 | -0.67 | 0.5214 | 0.7821 | -1.13 | 0.2656 | 0.7824 | -2.28 | 0.0262* | 0.1572 |
| Betweenness Centrality | L IPL ^a^ | 1.36 | 0.1740 | 0.4176 | 0.95 | 0.3454 | 0.6322 | 0.49 | 0.6370 | 0.9555 | 1.41 | 0.1696 | 0.4469 |
|  | L LC | 0.48 | 0.6244 | 0.7834 | 1.6 | 0.1160 | 0.348 | -0.2 | 0.8476 | 0.9564 | 0.95 | 0.3494 | 0.5133 |
|  | L cMTG ^a^ | -1.62 | 0.1046 | 0.4176 | -1.71 | 0.0676 | 0.348 | 1.59 | 0.1242 | 0.4968 | -0.99 | 0.3252 | 0.5133 |
|  | L pMTG ^a^ | 2.7 | 0.0104* | 0.0624 | 1.61 | 0.1132 | 0.348 | 1.58 | 0.1182 | 0.4968 | 3.01 | 0.0042** | 0.0504 |
|  | R aIPL | -1.2 | 0.2120 | 0.424 | -1.55 | 0.1158 | 0.348 | -0.28 | 0.7872 | 0.9564 | -1.53 | 0.1452 | 0.4469 |
|  | R LC | 1.45 | 0.1584 | 0.4176 | 0.35 | 0.7220 | 0.7876 | -0.1 | 0.9222 | 0.9564 | 0.89 | 0.3850 | 0.5133 |
|  | R STG ^c^ | -0.74 | 0.4838 | 0.7834 | -1.25 | 0.2210 | 0.5304 | -0.8 | 0.4424 | 0.8174 | -1.36 | 0.1862 | 0.4469 |
|  | VTA | -0.11 | 0.9116 | 0.9282 | -0.46 | 0.6464 | 0.7876 | 0.98 | 0.3260 | 0.7824 | 0.13 | 0.8978 | 0.8978 |
| **TAS:** Toronto Alexithymia Scale; **DIF:** Difficulty Identifying Feelings; **DDF:** Difficulty Describing Feelings; **EOT:** Externally Oriented Thinking; **L:** Left; **R:** Right; **B:** Bilateral; **vmPFC:** ventromedial Prefrontal Cortex; **DR:** Dorsal Raphe; **dmPFC:** dorsomedial Prefrontal Cortex; **IPL:** Inferior Parietal Lobule; **aIPL:** anterior IPL; **cMTG:** central Middle Temporal Gyrus; **pMTG:** posterior MTG; **LC:** Locus Coeruleus; **STG**: Superior Temporal Gyrus; **VTA:** Ventral Tegmental Area; **FDR:** False-Discovery Rate.  * p < 0.05; ** p < 0.01; *** p < 0.001  ^a^ Peak activity coordinates, MNI: **L IPL** (-46, -66, 46); **L cMTG** (-60, -61, 0); **L pMTG** (-64, -25, -5).  ^b^ Center of mass coordinates, MNI: **L aIPL** (-56, -38, 51). **L aIPL** includes two overlapping areas from the functional analysis: **L IPL / PCG** (Postcentral Gyrus) (peak activity coordinates: -55, -32, 57) and **L IPL** (peak activity coordinates: -57, -41, 50).  ^c^ This ROI pertains to the R STG0 subdivision as described in the work by Hoffman and Lambon-Ralph (2018). | | | | | | | | | | | | | |

**Supplementary Table 7.**

Associations between relevant findings and anxiety

| **Findings** | | **Trait-anxiety (STAI)** | | |
| --- | --- | --- | --- | --- |
|  |  | **t** | **p** | **FDR** |
| Modularity | | -1.035 | 0.3050 | 0.4270 |
| Clustering Coefficient | L pMTG ^a^ | -1.669 | 0.1010 | 0.1768 |
| Functional Connectivity | R STG ^b^ - L MTG | 2.689 | **0.0093**** | **0.0327*** |
|  | R LC - L IPL | -2.822 | **0.0065**** | **0.0327*** |
|  | DR - R IPL | -0.299 | 0.7660 | 0.7660 |
|  | DR - L IPL / PCG | -0.745 | 0.4590 | 0.5355 |
|  | R LC - B vmPFC | -1.826 | 0.0730 | 0.1703 |
| **L:** Left; **R**: Right; **B:** Bilateral; **MTG:** Middle Temporal Gyrus; **pMTG:** posterior MTG; **STG:** Superior Temporal Gyrus; **LC:** Locus Coeruleus; **IPL:** Inferior Parietal Lobule; **DR:** Dorsal Raphe; **PCG:** Postcentral Gyrus; **vmPFC:** ventromedial Prefrontal Cortex.  * p < 0.05; ** p < 0.01  ^a^ Peak activity coordinates, MNI: **L pMTG** (-64, -25, -5).  ^b^ This ROI pertains to the R STG0 subdivision as described in the work by Hoffman and Lambon-Ralph (2018). | | | | |

**Supplementary Table 8.**

Associations between relevant findings and suppression strategy

| **Findings** | | **Suppression (ERQ)** | | |
| --- | --- | --- | --- | --- |
|  |  | **t** | **p** | **FDR** |
| Modularity | | 0.046 | 0.9630 | 0.9700 |
| Clustering Coefficient | L pMTG ^a^ | -0.038 | 0.9700 | 0.9700 |
| Functional Connectivity | DR - L IPL | -1.225 | 0.2260 | 0.6027 |
|  | DR - L MTG | -0.945 | 0.3480 | 0.6960 |
|  | L LC - L dmPFC | -3.303 | **0.0016**** | **0.0131*** |
|  | DR - R IPL | -0.246 | 0.8070 | 0.9700 |
|  | DR - IPL / PCG | -1.249 | 0.2170 | 0.6027 |
|  | R LC - B vmPFC | -0.201 | 0.8410 | 0.9700 |
| **L:** Left; **R**: Right; **B:** Bilateral; **MTG:** Middle Temporal Gyrus; **pMTG:** posterior MTG; **DR:** Dorsal Raphe; **IPL:** Inferior Parietal Lobule; **LC:** Locus Coeruleus; **dmPFC: d**orsomedial Prefrontal Cortex; **PCG:** Postcentral Gyrus; **vmPFC:** ventromedial Prefrontal Cortex.  * p < 0.05; ** p < 0.01  ^a^ Peak activity coordinates, MNI: **L pMTG** (-64, -25, -5). | | | | |
